# Small-molecule CBLB inhibitor abolishes EGFR ubiquitination, reduces receptor endocytosis and diminishes cell motility signaling

**DOI:** 10.64898/2026.02.24.707836

**Authors:** Itziar Pinilla-Macua, Ratul Mukerji, Frederick Cohen, Alexander Sorkin

## Abstract

Endocytosis of the epidermal growth factor receptor (EGFR) is considered a key regulator of the receptor signaling activity. However, the molecular mechanisms underlying EGFR endocytosis are incompletely understood. Although ligand-induced ubiquitination of EGFR is known to promote its endocytic trafficking, the importance of EGFR ubiquitination in clathrin-mediated endocytosis, the primary physiological route of EGFR internalization, remains debated, and the relative contributions of ubiquitination-dependent and - independent mechanisms are not defined. Hence, we used NX-1013, a novel small-molecule inhibitor of the CBLB E3 ubiquitin ligase, to dissect the role of EGFR ubiquitination in its endocytic trafficking and signaling. Strikingly, brief treatment with NX-1013 completely abolished EGF-induced EGFR ubiquitination, demonstrating that this process is exclusively mediated by the closely related CBLB and CBL ligases. NX-1013 inhibited clathrin-mediated internalization of activated EGFR by 60–70%. The remaining, ubiquitination-independent internalization required EGFR kinase activity, was highly clathrin-dependent, and was significantly impaired by depletion of the AP-2 clathrin adaptor complex. Interestingly, inhibition of CBLs and EGFR endocytosis by NX-1013 did not affect major downstream signaling pathways in human oral squamous cell carcinoma cells, with the exception of Rac1 activation and EGFR-dependent cell migration, both of which were suppressed.

**Significance Statement:** CBL E3 ubiquitin ligases mediate ubiquitin conjugation of EGFR but their functional contributions to EGFR endocytic trafficking and signaling remain poorly defined. Here, we describe a newly developed small-molecule inhibitor of CBL proteins that potently blocks EGFR ubiquitination. This tool allowed us to dissect ubiquitination-dependent versus - independent components of the clathrin-mediated endocytosis and ligand-induced downregulation of EGFR. Strikingly, while inhibition of CBLs suppressed EGFR-driven cell motility signaling, it spared other major downstream pathways in EGFR-dependent human oral squamous cell carcinoma cells. These findings establish acute inhibition of CBLs as a powerful approach to interrogate ubiquitin-mediated receptor regulation and highlight its potential for therapeutic targeting of cancer cell migration.

## INTRODUCTION

The epidermal growth factor (EGF) receptor (EGFR) is the most extensively studied member of the receptor tyrosine kinase (RTK) family. EGFR is essential for mammalian development, wound healing and adult tissue homeostasis (1, 2). Mutations and overexpression of EGFR are common in a variety of cancers, making it a key prognostic marker and therapeutic target (3). The EGFR signaling system has served as a paradigm for elucidating the biology of other RTKs and signaling receptors in general. Ligand binding to EGFR at the cell surface results in activation of the tyrosine kinase in the cytoplasmic domain of the receptor leading to receptor autophosphorylation and phosphorylation of downstream signaling proteins. These events initiate multiple signal transduction processes that regulate cell proliferation, differentiation, metabolism, and motility (4). EGFR has also been a principal model for studying receptor-mediated endocytosis. Upon activation, EGFR is rapidly internalized and sorted in endosomes for lysosomal degradation (5). These endocytic and post-endocytic processes play crucial roles in modulating EGFR signaling (6, 7). However, the physiological mechanisms of EGFR endocytosis remain incompletely understood, as is the case for other RTKs.

Ligand-induced receptor ubiquitination is the key sorting signal during the endocytic trafficking of EGFR and several other RTKs [reviewed in (8)]. Ubiquitination was shown to be one of the redundant mechanisms of clathrin-mediated endocytosis (CME) of EGFR, the main physiological pathway of EGFR internalization (9, 10). However, the role of EGFR ubiquitination in the EGFR CME has been questioned because the CME of EGFR was observed under conditions when EGFR ubiquitination was negligible or undetectable (11–13). On the other hand, EGFR polyubiquitination was shown to be essential for the efficient post-endocytic sorting of internalized receptors to the lysosomal degradation pathway (14). EGFR ubiquitination is primarily mediated by Casitas B-lineage lymphoma (CBL) E3 ubiquitin ligases (15), although other ligases have been additionally implicated (16). In mammals, three CBL isoforms are expressed: CBL, CBLB, and CBLC. CBLs have a highly homologous amino-terminal tyrosine kinase binding (TKB) domain, mediating binding to phosphotyrosines of EGFR and other RTKs, a linker helix region (LHR), and a RING-type Zn finger domain that recruits an E2 ubiquitin (Ub)-conjugating enzyme. CBL and CBLB contain a variable unfolded proline-rich region that serves for binding of SH3 domain containing proteins, mediating indirect binding to RTKs, and a C-terminal ubiquitin associated domain [reviewed in (17, 18)]. CBLC lacks most of the C-terminal region, although it can bind to RTKs via its TKB domain. However, CBLC knockout mice exhibited normal development and EGFR signaling, and it is thought that CBLC is not directly involved in RTK ubiquitination [(11, 19, 20), reviewed in (21)].

Previous studies of the role of CBLs in EGFR ubiquitination and trafficking used dominant-negative CBL mutants, RNAi, or knockouts. These genetic approaches variably reduced EGFR ubiquitination, endocytosis and degradation, depending on cell type, experimental assays and compensatory mechanisms (9, 11, 22–26). To more directly probe the contribution of CBLs to EGFR ubiquitination and trafficking we took advantage of compounds emerging from a high-throughput screening campaign designed to identify inhibitors of CBLB for use as immunotherapeutics (27). A series of potent small molecule triazole-based inhibitors were identified and optimized to yield NX-1607, which is currently being evaluated in clinical trials of advanced malignancies (28, 29) (clinical trials.gov ID: NCT05107674). Structural studies confirmed that this triazole-based series binds to the TKB domain and LHR of CBLB, acting as an intramolecular glue to lock the protein in a closed and inactive conformation, and predicted that these compounds can also inhibit CBL (27). While the ability of NX-1607 to activate immune cells has been demonstrated (28, 29), the cell-autonomous effects of this series of compounds on solid tumor cells are not well characterized. In this study, we used NX-1013, a close analog of the clinical compound NX-1607 (NCT05107674) with similar properties and activity, to investigate its effects on EGFR ubiquitination, trafficking, and signaling in human oral squamous cell carcinoma cells, which rely on EGFR for growth. Remarkably, short treatment with NX-1013 abolished EGF-induced EGFR ubiquitination, indicating that CBL proteins are uniquely responsible for EGFR ubiquitination. NX-1013 also strongly inhibited EGFR CME and endosomal sorting, although a substantial residual endocytosis and ligand-induced receptor degradation persisted in the absence of receptor ubiquitination. Acute inhibition of EGFR ubiquitination allowed us to demonstrate that ubiquitination-independent endocytosis of endogenous EGFR requires its tyrosine kinase activity, clathrin and the clathrin adaptor complex AP-2.

## RESULTS AND DISCUSSION

### Mechanism of action and biochemical characterization of NX-1013

CBL proteins exhibit a closed conformation that is incapable of binding an Ub-loaded E2 Ub-conjugating enzymes (Fig. 1A). Substrate binding triggers a conformational rearrangement to an open state that allows phosphorylation of a key tyrosine residue (Y363 in CBLB) which then allows Ub-E2 binding and subsequent Ub transfer to substrate proteins (30, 31). NX-1013 binds to the closed conformation and functions as an intramolecular glue, shifting the conformational equilibrium toward the closed state (Fig. 1B). Biophysical characterization by surface plasmon resonance (SPR) indicated high-affinity binding of NX-1013 to the closed conformations of both CBLB and CBL. The data were best fit to a 1:1 binding model, yielding K_d_ of 0.87 ± 0.09 nM for CBLB and 3.4 ± 0.6 nM for CBL (Fig. 1C). Orthogonal confirmation of binding was provided by a TR-FRET assay employing a Bodipy-Fluorescein-labeled analog of NX-1013 (Fig. 1D). In this assay, NX-1013 displayed IC_50_ values of 0.51 ± 0.27 nM and 1.7 ± 1.3 nM for CBLB and CBL respectively. Screening of a close analog of NX-1013 (compound 7) against a panel of 55 enzymes, receptors, transporters, and channels at 10 μM showed that most targets were not affected and no target was inhibited by more than 40%, indicating that NX-1013-like compounds are not promiscuous inhibitors (27).

**Figure 1.**
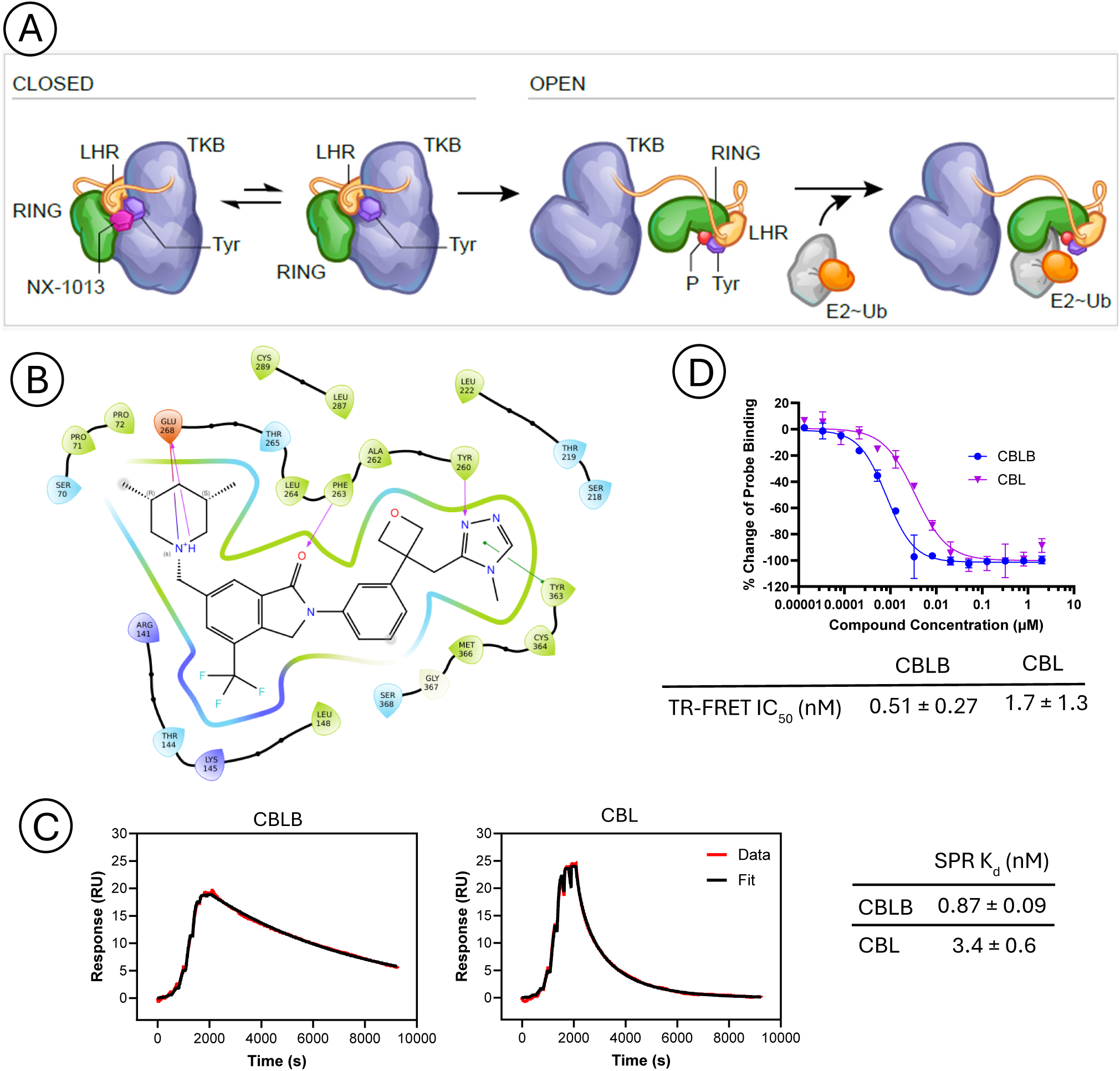
Mechanism and characterization of NX-1013. **(A)** CBL proteins are in equilibrium between open and closed states. NX-1013 binds to the closed state, functioning as an intramolecular glue between LHR and TKB domains to stabilize the closed conformation, preventing phosphorylation and activation. **(B)** Chemical structure and ligand interaction diagram. A model of NX-1013 binding to CBLB was created from co-crystal structure of a close analog (PDB9XZD). Key protein-ligand interactions include a hydrogen bond between N2 of the triazole ring and the phenol of Tyr260; a *pi-pi* stacking between the triazole and Tyr363 as well as a salt bridge between the charged piperidine and side chain of Glu268. **(C)** Surface Plasmon Resonance (SPR) analysis demonstrated high-affinity binding of NX-1013 to the closed conformations of both CBLB and CBL. The sensorgrams were best fit to a 1:1 binding model. **(D)** Orthogonal confirmation of binding. NX-1013 shows a dose-dependent signal decrease in the TR-FRET probe displacement assay, resulting in IC_50_ values of 0.51 ± 0.27 nM and 1.7 ± 1.3 nM for CBLB and CBL respectively.

### NX-1013 is a highly effective inhibitor of EGFR ubiquitination

To examine the effects of NX-1013 on EGFR ubiquitination, human oral squamous cell carcinoma (HSC3) cells were used because they express a relatively high level of CBLB (26) and are growth-dependent on the EGFR kinase activity (32). Immunoprecipitation of EGFR from HSC3 cells stimulated with EGF for 10 min reveals strong EGF-concentration dependent receptor ubiquitination (Fig. 2A-B). Remarkably, 100 nM NX-1013 decreased EGF-induced EGFR ubiquitination detectable by western blotting to the level of ubiquitination in unstimulated cells (Fig. 2A-B). When only a few receptors were activated (1 ng/ml EGF), NX-1013 partially reduced CBLB association with EGFR (Fig. 2A, C). A trend of a reduced CBLB co-precipitation with EGFR activated with 10-100 ng/ml EGF in the presence of NX-1013 was observed but this reduction was not statistically significant. The effects of NX-1013 on CBL association with the receptor were also not significant, although a trend for decreased CBL co-immunoprecipitation, when using 1 μM NX-1013 and 1 ng/ml EGF, was observed (Fig. 2A and D). The results of co-immunoprecipitation experiments imply that while structural studies predict that binding of NX-1013-like compounds does not affect the phosphotyrosine binding interface of the TKB domain of CBLB (27), this prediction must be experimentally tested.

**Figure 2.**
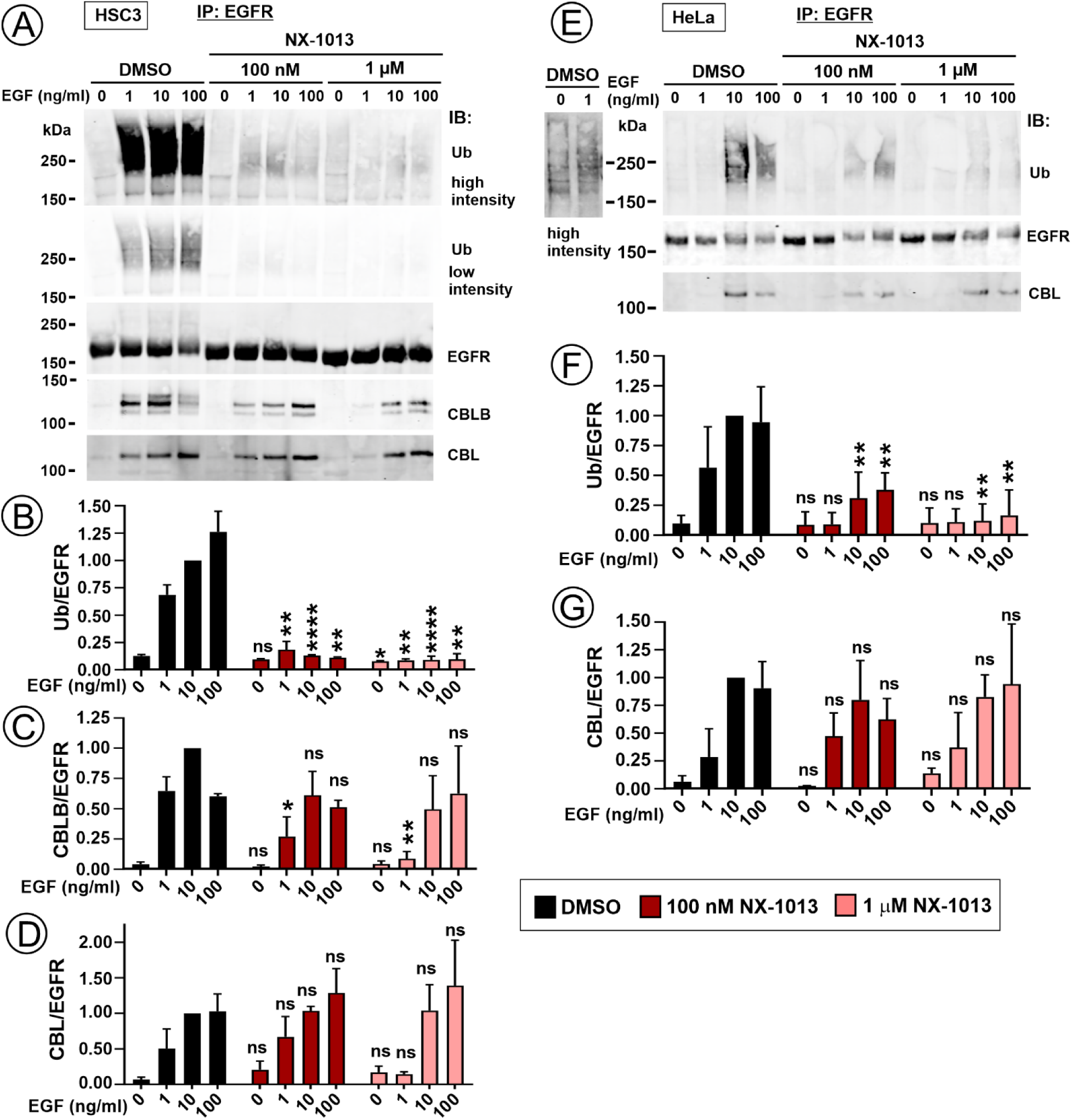
Effect of NX-1013 on EGFR ubiquitination and association with CBLs. (**A**) HSC3 cells were preincubated with DMSO (vehicle), 100 nM or 1 μM NX-1013 for 1 hr and stimulated with 1-100 ng/ml EGF for 10 min at 37°C. EGFR was immunoprecipitated from lysates, and immunoprecipitates were probed by western blotting with antibodies to ubiquitin (Ub), EGFR, CBL and CBLB. Representative images are shown. (**B-D**) Quantification of EGFR ubiquitination (***B***), amount of co-immunoprecipitated CBLB (***C***) and CBL (***D***) from western blots exemplified in ***A***. Bars represent mean values (± SEM; n=3 independent experiments) of the ratio of ubiquitin, CBL or CBLB to EGFR immunoreactivities normalized to this ratio in DMSO-treated cells stimulated with 10ng/ml EGF in each individual experiment. **(E)** HeLa cells were preincubated with DMSO, 100 nM or 1 mM NX-1013 for 1 hr and stimulated with 1-100 ng/ml EGF for 10 min at 37°C. EGFR was immunoprecipitated from lysates, and immunoprecipitates were probed by western blotting with antibodies to Ub, EGFR and CBL. Representative images are shown. (**F-G**) Quantification of EGFR ubiquitination (***F***) and amount of co-immunoprecipitated CBL (***G***) from western blots exemplified in ***E***. Bars represent mean values (± SEM; n=3 independent experiments) of the ratio of ubiquitin or CBL to EGFR immunoreactivities normalized to the ratio in DMSO-treated cells stimulated with 10ng/ml EGF in each individual experiment. (**B**, **C**, **D**, **F** and **G**) Differences between DMSO- and NX-1013-treated variants at corresponding EGF concentration were calculated using one-way ANOVA: *p<0.05; **p<0.01; ***p<0.001; ****p<0.0001; ns, not significant (p>0.05). Actual p values are summarized in Table S1. For differences of EGF stimulation versus unstimulated cells within group variants exposed to the same concentration of NX-1013, p values (not shown in the figure) were calculated using one-way ANOVA and are presented in Table S1.

Most cultured human cells of epithelial origin express very low levels of CBLB as compared to CBL, unlike HSC3 cells, which are the main experimental model in our studies. Therefore, we examined the effects of NX-1013 on EGFR ubiquitination in human cervical carcinoma HeLa cells expressing similar level of CBL as in HSC3 cells but ∼10-times less CBLB than CBL (26). HeLa cells also express an order of magnitude lower level of EGFR than do HSC3 cells. As expected, EGFR ubiquitination was at the margin of detection sensitivity by western blotting in HeLa cells treated with 1 ng/ml EGF (see “high intensity” inset in Fig. 2E). Receptor ubiquitination was highest in cells treated with a subsaturating EGF concentration (10 ng/ml) (Fig. 2E-F). NX-1013 strongly decreased EGF-induced EGFR ubiquitination in HeLa cells under all conditions although higher NX-1013 concentration (1 μM) was necessary for the maximal extent of the inhibition (Fig. 2E-F) as compared to HSC3 cells. NX-1013 did not affect co-immunoprecipitation of CBL with EGFR activated with high EGF concentrations (10-100 ng/ml). A statistically significant association of CBL with EGFR in cells treated with 1 ng/ml EGF was not detected because of a low signal-to-noise ratio. Likewise, because of a low expression level, we have not been able to detect CBLB co-immunoprecipitated with EGFR in HeLa cells.

EGF-induced EGFR ubiquitination involves direct and indirect binding of CBLs to the receptor, which requires receptor tyrosine phosphorylation and therefore its kinase activity. To rule out indirect effects of NX-1013 on the activation of EGFR and its downstream signaling, cell lysates were probed for active EGFR, extracellular signal-regulated kinase 1/2 (ERK1/2) and Akt. Figure S1 shows that NX-1013 did not affect EGFR phosphorylation at the major site, Y1068, or signaling through two major pathways under experimental conditions used in Figure 2 in both HSC3 and HeLa cells.

Overall, EGFR immunoprecipitation experiments presented in Figure 2 demonstrate that NX-1013 is a highly effective inhibitor of EGFR ubiquitination. A strong effect of NX-1013 on EGFR ubiquitination in HeLa cells suggests that CBL is also inhibited by this compound in cells, albeit with a lesser efficiency as compared to that towards CBLB, thus confirming our *in vitro* analyses (Fig. 1C-D). Together, these data, combined with the demonstrated high specificity of NX-1013–like compounds (27) and the uniqueness of the compound-interacting molecular determinants in TKB and LHR domains of CBLB and CBL across the entire E3 ubiquitin ligase family (Fig. 1B and ref. (27)), support the conclusion that CBL proteins are the only physiological E3 Ub ligases for EGFR. NX-1013 has shown greater effectiveness in inhibiting EGFR ubiquitination compared to CBL knockdowns or dominant-negative mutant overexpression in HSC3 cells (26). There may be at least two possible explanations for these observations. First, the latter approaches may preserve small amounts of active CBL and/or CBLB which can be sufficient for residual receptor ubiquitination. Second, in the absence of CBL and CBLB activities, adaptive changes may result in CBLC or other E3 ligases acting to catalyze EGFR ubiquitination. In fact, CBLC is expressed at a high level in HSC3 cells (https://cansar.ai/) whereas it is undetectable in HeLa cells (11).

### NX-1013 strongly inhibits endocytic trafficking of EGFR

The high efficacy of NX-1013 in blocking EGFR ubiquitination provides a unique opportunity for defining the role of ubiquitination in receptor endocytosis in cells expressing endogenous EGFR while avoiding genetic manipulations and adaptive changes. Thus, we monitored the effect of NX-1013 on trafficking of EGF conjugated to rhodamine (EGF-Rh; 10 ng/ml) in HSC3 cells using time-lapse, three-dimensional (3D) imaging by a Lattice LightSheet microscopy system. In vehicle-treated cells, EGF-Rh rapidly accumulated in numerous vesicular compartments, presumably endosomes, which displayed flickering motion motility and tended to accumulate in the perinuclear area (Fig. 3 and VideoS1). By contrast, the appearance of EGF-Rh in vesicles was delayed and the density of vesicles was much lower in NX-1013-treated than in control cells (Fig. 3 and VideoS2). Cell-edge EGF-Rh labeling typical of the surface localization was faint after 10-15 min in control cells whereas such labeling, including numerous immobile clusters of EGF-Rh, was apparent in NX-1013-treated cells during at least 30 min of time-lapse imaging (VideoS2).

**Figure 3.**
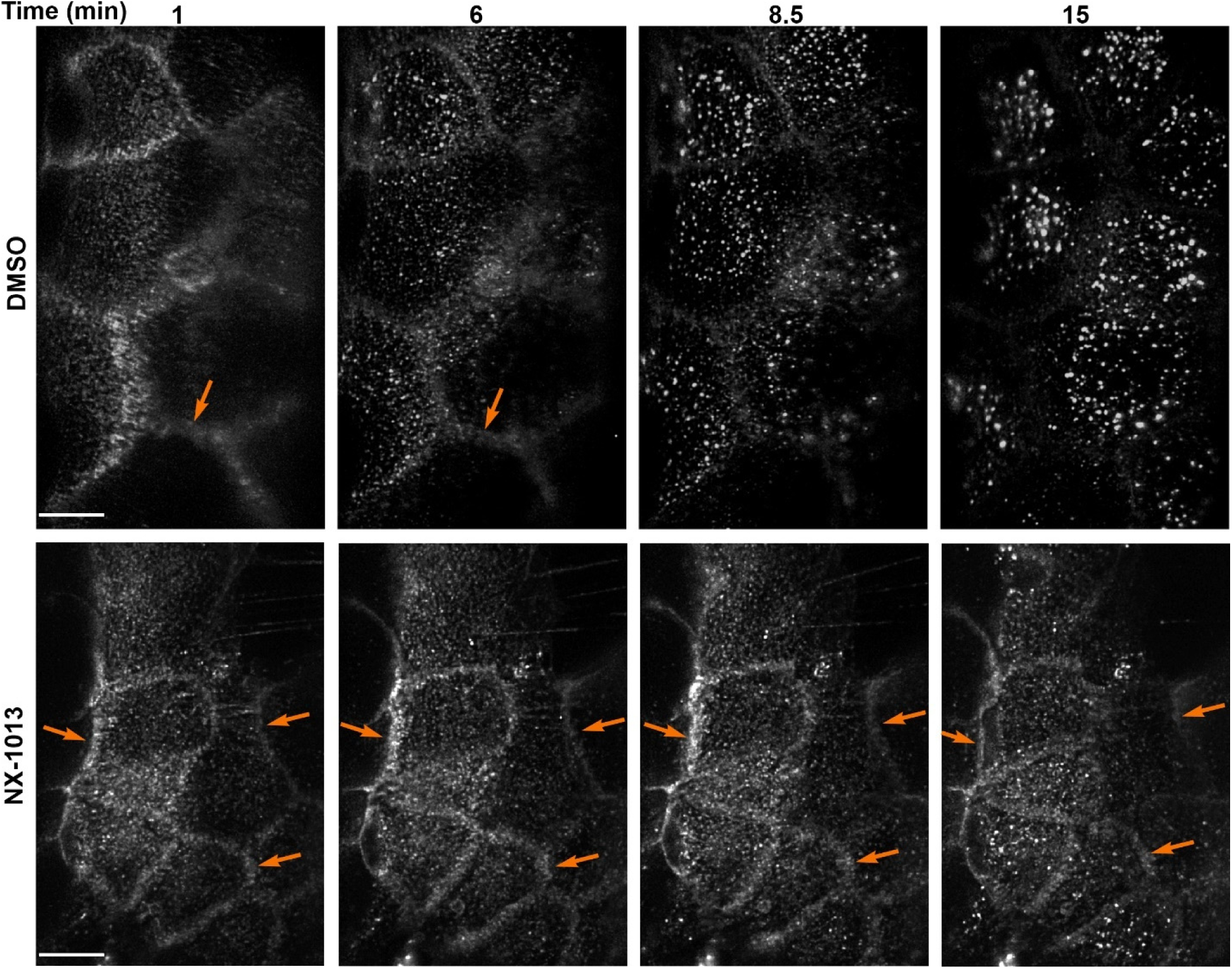
Live-cell imaging reveals delayed EGFR endocytosis in HSC3 cells treated with NX-1013. Cells were pre-incubated with DMSO or 100 nM NX-1013 and then 3D images were acquired using the LLS system every 30 seconds starting 1 min after injection of EGF-Rh in the presence of DMSO or NX-1013 at 37°C. Maximum intensity projections of z-stack of x-y images of selected time points from time-lapse sequences are shown. Scale bars are 10 μm. Arrows point to examples of EGF-Rh localization at the cell surface. See corresponding Videos S1 and S2 in *Supplemental Information*. The experiment is representative of 4 (vehicle) and 15 (NX-1013) time-lapse imaging experiments where similar kinetics of endocytosis were observed.

Immunofluorescence microscopy confirms the slowed endocytosis of non-ubiquitinated EGFR in HSC3 cells (Fig. 4). In control cells, a large fraction of EGF-Rh was found in the perinuclear early/sorting endosomes labeled with EEA1 after 10 min of EGF-Rh (4 ng/ml) stimulation (Fig. 4A and C). The fraction of EGF-Rh located in EEA1-labeled endosomes was smaller, and the endosomes were distributed throughout the cell in NX-1013-treated cells (Fig. 4B-C). The plasma membrane localization of EGF-Rh remained clearly visible after 1 hour of continuous EGF-Rh endocytosis in NX-1013-treated cells (Fig. 4B). As previously shown, the sorting of EGFR to late endosomes and ligand-induced EGFR degradation are relatively slow in HSC3 cells (33). Consistently, a rather small fraction of EGF-Rh was found in LAMP1 labeled compartments (late endosomes and lysosomes) in vehicle-treated cells after 1-hour incubation with EGF-Rh (Fig. 4A and C). In NX-1013-treated cells, no increase in EGF-Rh/LAMP1 colocalization above the basal level of an apparent fluorescence overlap was detected (Fig. 4C). To test whether inhibition of EGF-Rh endocytosis by NX-1013 reflects inhibition of EGFR endocytosis, localization of EGFR was examined by immunofluorescence labeling before and after cell stimulation with EGF-Rh. NX-1013 did not affect the amount or distribution of EGFR at the cell surface of unstimulated cells (Fig. S2A). While 4 ng/ml EGF-Rh resulted in endocytosis of only a small fraction of EGFR in HSC3 cells, internalized EGFR was highly colocalized with EGF-Rh in endosomes (Fig. S2B-C). NX-1013 significantly decreased endocytic accumulation of EGFR in EEA1-labeled endosomes (Fig. S2B-D). Together, live-cell imaging and immunofluorescence analysis demonstrate that EGFR endocytosis and post-endocytic trafficking of non-ubiquitinated EGFR are significantly slower than that of ubiquitinated EGFR, although the ubiquitination-independent endocytic trafficking of the receptor is still substantial.

**Figure 4.**
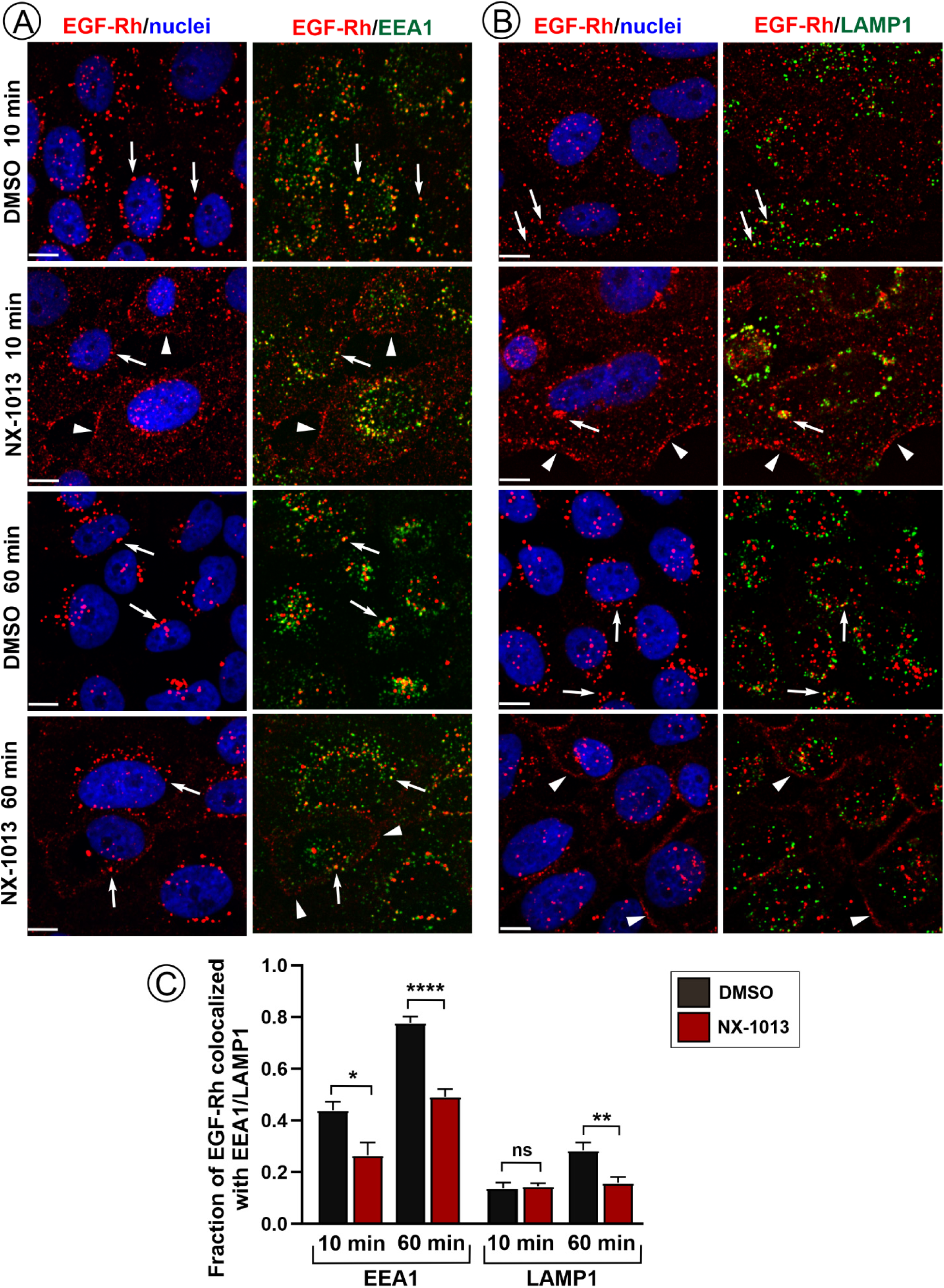
Delayed endocytic trafficking to early and late endosomes in HSC3 cells treated with NX-1013. Cells were pre-incubated with DMSO or 100 nM NX-1013 and then incubated with 4 ng/ml EGF-Rh for 10 or 60 min at 37°C in presence or absence of the inhibitor, fixed, permeabilized and labeled with EEA1 (**A**) or LAMP1 (**B**) antibodies. Imaging was through the 640 nm (*green*, EEA1/LAMP1), 561 nm (*red*, EGF-Rh) and 405 nm (*blue*, nuclei) channels. Maximum intensity projections of z-stacks are presented. Fluorescence intensity scales are identical for all images. White arrows mark examples of EGF-Rh puncta colocalized with EEA1 or LAMP1. Arrowheads point to the plasma membrane. Scale bars, 10 μm. (**C**) Quantifications of the fraction of EGF-Rh co-localized with EEA1 or LAMP1 of total cell-associated EGF-Rh in images exemplified in (***A***) and (***B***). Bar graph represents mean values (± SEM; n=4-6 FOVs) of the fraction of EGF-Rh co-localized with EEA1 or LAMP1. Unpaired two-tailed Student T-test between vehicle and NX-1013-treated cells for the same incubation times was used to calculate p values. *p<0.05 (p=0.019); **p<0.01 (p=0.0095); ****p<0.0001; ns, not significant (p>0.05).

To quantitatively characterize NX-1013 effects on the CME of EGFR, the major physiological pathway of EGFR endocytosis, low concentration of ^125^I-EGF (1 ng/ml) and short incubation times were used. These conditions favor EGFR internalization through the saturable CME pathway and minimize the contribution of EGF-receptor complex recycling in the overall ligand uptake (5). In HSC3 cells, a maximum inhibition of internalization (∼50-55%) was achieved by 100 nM NX-1013 (Fig. 5A-B). NX-1013 did not affect ^125^I-EGF binding to receptors, as measured using the equilibrium binding assay at 4°C (Fig. S3A), supporting the notion that the activity of CBLs does not affect dormant EGFR (Fig. S2A). NX-1013 significantly reduced internalization (by 60%) in cells preincubated with ^125^I-EGF at 4°C and then chased at 37°C (Fig. S3B). EGFR is capable of efficient accumulation in clathrin coated pits in the absence of endocytosis when cells are incubated with EGF for ∼1-hr at 4°C, conditions of the ligand-receptor binding equilibrium (10). Therefore, inhibition of ^125^I-EGF internalization by NX-1013 after ^125^I-EGF pre-binding to receptors at 4°C (Fig. S3B) is indicative of a reduced efficiency of the recruitment of non-ubiquitinated EGFR into coated pits. This hypothesis is consistent with the 2-fold decrease in coated pit recruitment of the EGFR mutant lacking 21 lysines after cell incubation with EGF at 4°C (10). Observations in Fig. S3B contrast with the lack of an effect of CBL/CBLB double-knockout on EGFR internalization in mouse embryonic fibroblasts (MEFs), which was observed using the same ^125^I-EGF internalization protocol (13). These differences are likely to be due to a low EGFR expression in MEFs and consequently a low number of cell-surface ^125^I-EGF:receptor complexes which could have been well below the capacity of the ubiquitination-independent pathway in these cells, and/or the result of adaptive compensatory changes.

**Figure 5.**
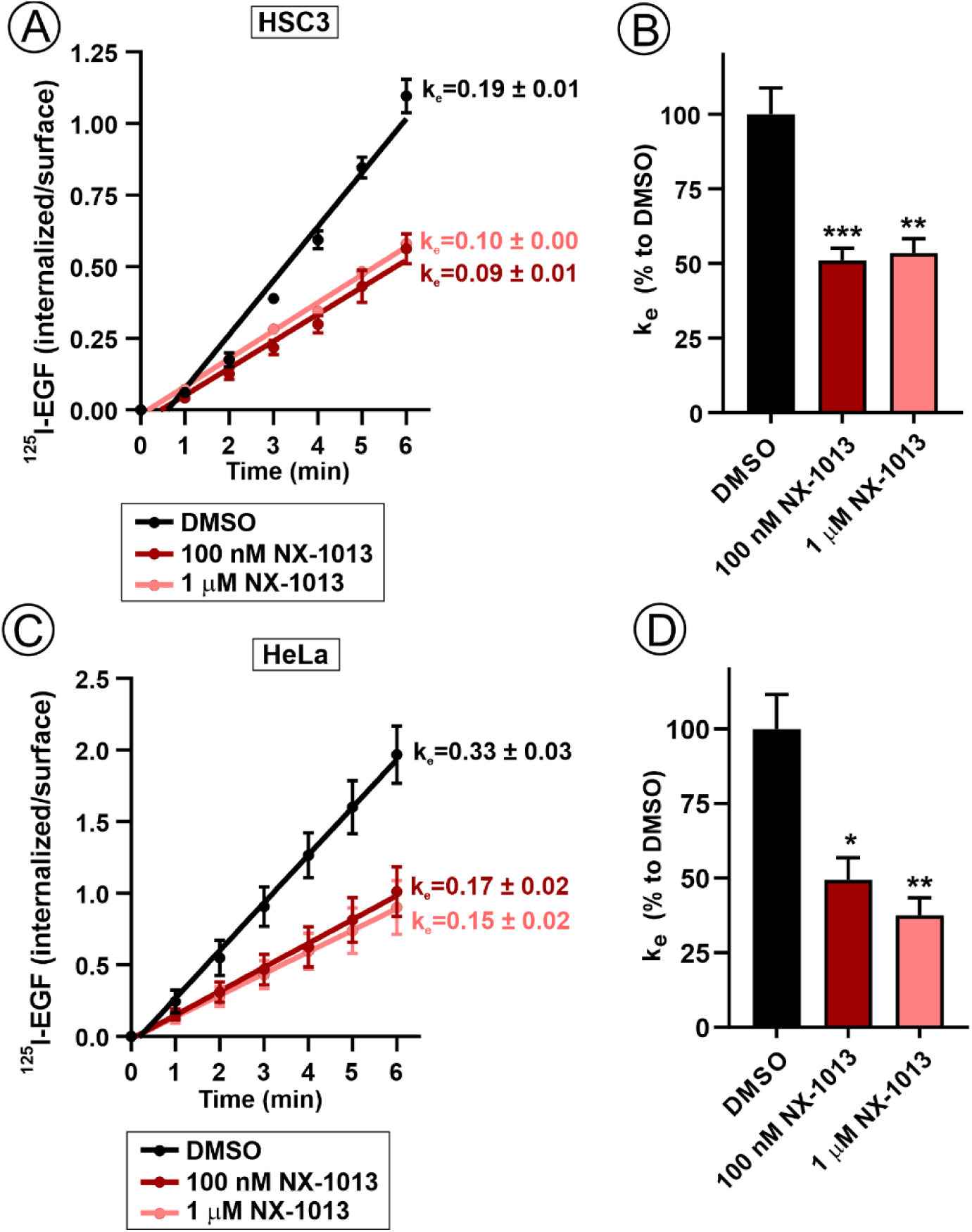
NX-1013 inhibits CME of EGFR. (**A-C**) HSC3 (**A**) or HeLa cells (**C**) were preincubated with DMSO (vehicle), 100 nM or 1 μM NX-1013 for 1 hr and then incubated with ^125^I-EGF (1 ng/ml) for 1-6 min. Internalized/surface ^125^I-EGF ratio and internalization rate constant k_e_ values (min^-1^) were quantitated as described in “Methods”. Graphs show mean values of internalization ratio for each timepoint (± SEM; n = 1-7 in ***A*** and n = 3-5 independent experiments in ***C***). (**B-D**) Mean values (± SEM) of k_e_ values were calculated from multiple independent experiments with HSC3 (n = 3-8) (**B**) and HeLa cells (n = 3-5) (**D**) as described in “Methods” and expressed as percent to mean values of k_e_ in vehicle-treated cells in individual experiments. P values of NX-1013-treated against DMSO-treated are calculated by one-way ANOVA. *p<0.05 (p = 0.021); **p< 0.01 (p = 0.0074 in **B** and p = 0.0032 in **D**); ***p<0.001 (p = 0.0008).

Analysis of the EGFR CME using EGF-Rh endocytosis assays presented in Figures 3 and 4 is not technically feasible and practical in HeLa cells due to a low level of EGFR and a significant clathrin-independent endocytosis at concentrations of EGF higher than 1-2 ng/ml (34). Nevertheless, ^125^I-EGF internalization assay demonstrated that the maximum extent of ^125^I-EGF CME inhibition by NX-1013 in HeLa cells is similar to that in HSC3 cells (Fig. 5C-D). Ubiquitination of EGFR stimulated with low concentrations of EGF (1-1.5 ng/ml EGF) has been technically challenging to detect by immunoblotting in HeLa cells [Fig. 2; also (11)], and therefore, the importance of EGFR ubiquitination for its CME was questioned (11, 12). The data in Figure 5 show that the CME of EGFR in these cells, as in HSC3 cells, is dependent on the E3 ubiquitin ligase activity of CBLs, and therefore, on EGFR ubiquitination. Previously, we demonstrated that mutations of major ubiquitination sites in EGFR resulted in ∼50% decrease in the internalization rate of the recombinant EGFR expressed in cells lacking endogenous EGFR (9, 10). The data in Figures 5 and S3 demonstrate a similar or a stronger (in some experiments) inhibition of the internalization of endogenous EGFR in acute-treatment experiments. Considering that constitutive internalization [k_e_ ∼ 0.01-0.04/min; (35)] contributes to the total uptake of ^125^I-EGF, the data in Figure 5 indicate that NX-1013 reduces the internalization rate constant of the EGFR CME by up to 60-70% in HSC3 and HeLa cells.

### Ubiquitination-independent internalization of EGFR

The effectiveness of NX-1013 in blocking ubiquitination of endogenous EGFR allowed us to examine the mechanisms of EGFR endocytosis that are independent of its ubiquitination. First, we found that Erlotinib, a small-molecule inhibitor of the EGFR kinase activity, slows internalization of ^125^I-EGF (1 ng/ml) in the presence of NX-1013 in HSC3 cells down to the level of constitutive internalization (k_e_ ∼ 0.03 min^-1^; Fig. S4A), demonstrating that EGFR kinase activity is essential for the ubiquitination-independent endocytosis. Second, the cells were transfected with the well-characterized siRNA duplex targeting clathrin heavy chain (CHC) to determine whether ubiquitination-independent endocytosis occurs via clathrin coated pits. CHC was depleted on average by 93% in HSC3 cells, which resulted in a substantial decrease of ^125^I-EGF (1 ng/ml) internalization rate in these cells untreated or treated with NX-1013 (k_e_=0.06 ± 0.01 min^-1^) (Fig. 6A). Interestingly, CHC knockdown significantly decreased internalization of ^125^I-EGF used in higher concentrations (4 and 10 ng/ml) in control and NX-1013-treated HSC3 cells (Fig. S3C-D), which is indicative of a large saturating capacity of the CME pathway for EGFR in these cells. The latter observations support the validity of microscopy observations in Figures 4 and 3 made using 4 and 10 ng/ml EGF, respectively. A highly efficient depletion of CHC in HeLa cells (>99%) led to the strongest inhibition of internalization in the absence or presence of NX-1013, down to the rate comparable to that of the constitutive EGFR internalization (k_e_=0.03±0.01 min^-1^) (Fig. 6C). Therefore, based on the internalization rate measurements in HeLa cells where the highest extent of CHC depletion was achieved, we conclude that the CME is the major pathway of the ubiquitination-independent endocytosis of ligand-bound EGFR. However, the data in Figures 6 and S2 demonstrate that the relative contribution of clathrin-independent endocytosis of EGFR in the overall uptake of ^125^I-EGF is larger in HSC3 than in HeLa cells regardless of the presence or absence of NX-1013.

**Figure 6.**
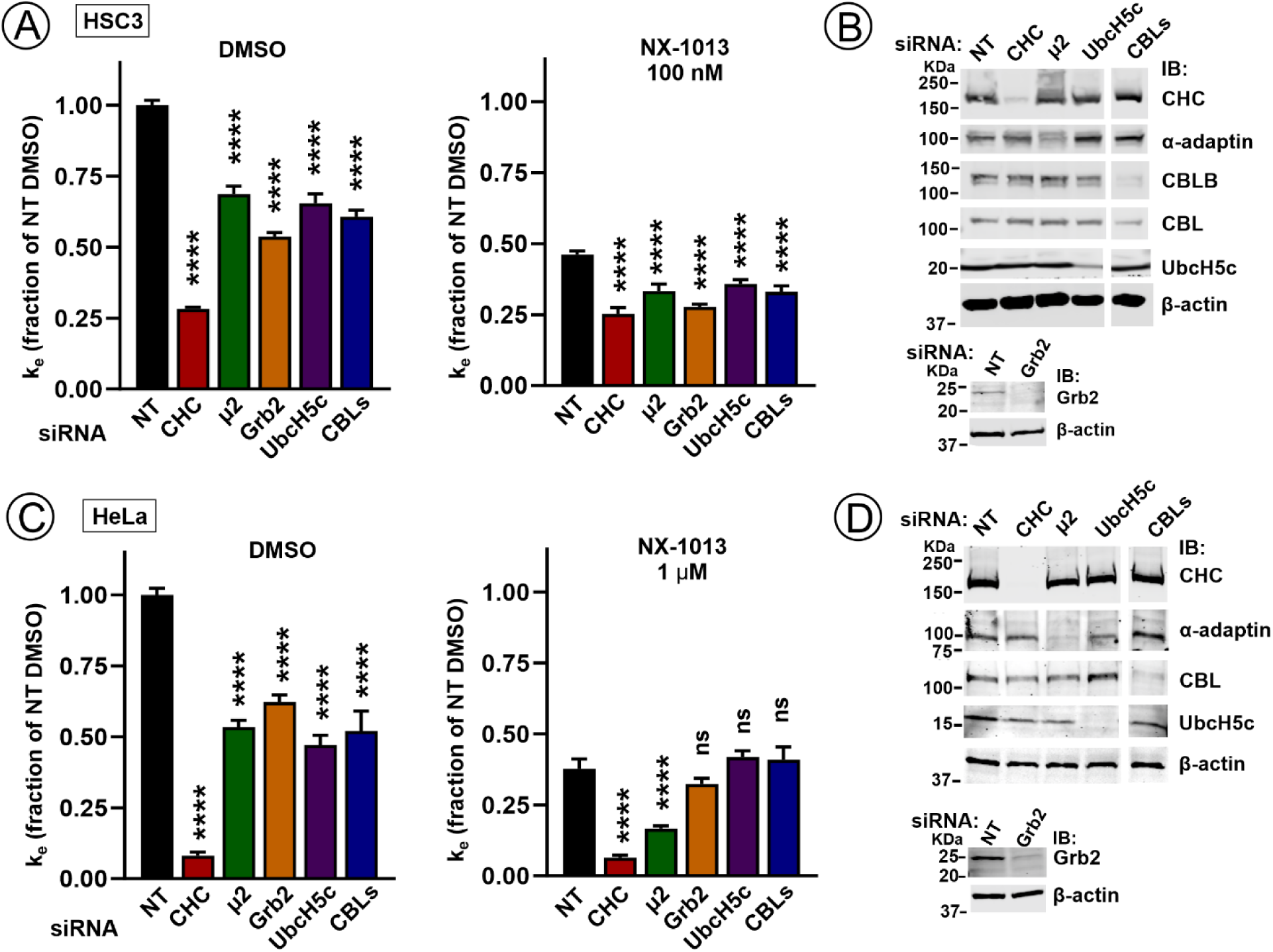
Ubiquitination-independent endocytosis of EGFR is clathrin-mediated. **(A)** HSC3 cells were transfected with non-targeting siRNA (NT) or siRNA targeting indicated proteins. Four days after transfection, the cells were preincubated with DMSO or 100 nM NX-1013 for 1 hr and then incubated with ^125^I-EGF (1 ng/ml) for 6 min. Bar graphs represent mean k_e_ values (± SEMs, n = 3-6 independent experiments, triplicates in each) normalized to k_e_ values in siNT-transfected, DMSO-treated cells in each individual experiment. P values are calculated against siNT-transfected cells using one-way ANOVA. ****p<0.0001. **(B)** Representative western blots showing extents of protein depletion in experiments presented in (***A***). The panels are from the same western blot (except the panel with Grb2 siRNA). **(C)** HeLa cells were transfected with NT siRNA or siRNA targeting indicated proteins. For experiments, the cells were preincubated with DMSO or 1 μM NX-1013 for 1 hr and then incubated with ^125^I-EGF (1 ng/ml) for 6 min. Bar graphs represent mean k_e_ values (± SEMs, n = 3-6 independent experiments, triplicates in each) normalized to k_e_ values in NT-transfected, vehicle-treated cells in each individual experiment. P values are calculated against siNT-transfected cells using one-way ANOVA. ****p<0.0001; ns, not significant (p>0.05). **(D)** Representative western blots showing extents of protein depletion in experiments presented in (***C***). The panels are from the same western blot (except the panel with Grb2 siRNA).

To examine the mechanism of ubiquitination-independent CME of EGFR, clathrin adaptor protein complex AP-2, a major clathrin coat component, was depleted by siRNA targeting the μ2 subunit of AP-2. Knockdown of AP-2 partially inhibited total and ubiquitination-independent ^125^I-EGF internalization in both HSC3 and HeLa cells (Fig. 6A-B). A stronger effect of this knockdown in HeLa cells is likely due to higher efficiency of AP-2 depletion (Fig. 6C-D). Subsequently, additional RNAi experiments were performed to target proteins known to be involved in EGFR ubiquitination and endocytosis. Depletion of the Grb2 adaptor that mediates indirect binding of CBL to EGFR was previously shown to strongly inhibit EGFR CME (34, 36). As shown in Figure 6, ubiquitination-independent EGFR endocytosis was slightly inhibited by Grb2 knockdown in HSC3 but not in HeLa cells. Knockdown of UBCH5b/c, E2 ubiquitin-conjugating enzymes associated with CBL and mediating EGFR ubiquitination (37), also somewhat reduced EGFR internalization in NX-1013-treated HSC3 but not in HeLa cells. Simultaneous depletion of CBL and CBLB affected EGFR internalization in a manner similar to what was observed in the case of Grb2 or UbcH5b/c depletion (Fig. 6). The most plausible explanation for the effects of Grb2, UbcH5b/c and CBLs knockdowns on ubiquitination-independent EGFR internalization in HSC3 cells is the presence of a residual receptor ubiquitination in these cells exposed to NX-1013, which is below the detection threshold of immunoblotting. By contrast, in HeLa cells expressing a lower total level of CBLs and an order of magnitude lower amount of EGFR than do HSC3 cells, NX-1013 may achieve its maximal efficiency in blocking the activity of CBLs and EGFR ubiquitination. Considering the data in Figure 6A, ubiquitination-independent mechanisms appear to control only a small fraction of the total EGFR internalization flow through the CME pathway in HSC3 cells, and it is not apparent that AP-2 has a major role in these mechanisms. A higher efficiency of siRNA knockdowns in HeLa cells allows a clearer interpretation of RNAi experiments in these cells: ubiquitination-independent EGFR CME is reliant on AP-2 but not on the residual activity of the receptor ubiquitination system.

The dependence of EGFR CME on AP-2 has been previously demonstrated in several human immortalized cell lines using RNAi knockdowns (12, 38) and in MEFs derived from the AP-2 knockout (12). Importantly, the latter study showed that elimination of AP-2 results in a dramatically reduced number of clathrin coated structures per cell rather than implicating the internalization process involving the direct interaction of the receptor with AP-2. A relatively small component of EGFR CME involving the dileucine- and tyrosine-based AP-2 interaction motifs of EGFR was revealed in cells expressing non-ubiquitinated EGFR mutant (10). Furthermore, endocytosis of ligand-free EGFR induced by the activity of p38 MAP kinase was recently shown to be mediated by the receptor di-leucine-motif-mediated interaction with AP-2 (34). However, inhibition of p38 MAP kinase did not affect ^125^I-EGF internalization in control and NX-1013-treated HSC3 cells (Fig. S4B), confirming that this mechanism is operational only with inactive receptors. The possibility remains that Y^974^RAL, LL1000/1001 and three NPxY motifs in the carboxy-terminal domain of EGFR, all capable of direct or indirect interactions with AP-2 and clathrin, mediate ubiquitination-independent endocytosis of EGFR in a redundant fashion. To conclude, the precise molecular mechanisms of ubiquitination-independent CME of EGFR remain to be elucidated. NX-1013 has a strong potential to be an indispensable experimental tool in the future mechanistic studies of both ubiquitination-dependent and -independent CME of EGFR and other RTKs in cultured cells and *in vivo*.

### Effects of inhibiting CBLs on EGFR signaling

It is generally assumed that the inhibition of RTK endocytosis leads to upregulation of signaling from the plasma membrane through major signaling pathways such as RAS-ERK1/2 and PI3K-Akt [reviewed in (6, 7)]. Does the acute inhibition of CBL/CBLB Ub ligase activity and slowdown of EGFR endocytosis influence the intensity and dynamics of these signaling processes downstream of activated EGFR? To address this question, we monitored EGFR activity and signaling outcomes in HSC3 cells stimulated with 4 ng/ml EGF. As shown in Figure 7A-B, EGFR activation, measured using antibodies to pY1068, was not affected by NX-1013. Similarly, the time-course of the activation of ERK1/2 and Akt was not different in control and NX-1013-treated cells (Fig. 7A, C-D), indicating that in cells like HSC3, with a high level of EGFRs, even a substantial decrease of EGFR CME does not significantly impact these two signaling pathways. Low EGF concentrations activate only a small fraction of EGFRs in HSC3 cells and do not significantly down-regulate total cell-surface EGFR, which results in the maintenance of the level of active receptors at the cell surface within a similar range regardless of whether endocytosis is fully operational or partially inhibited.

**Figure 7.**
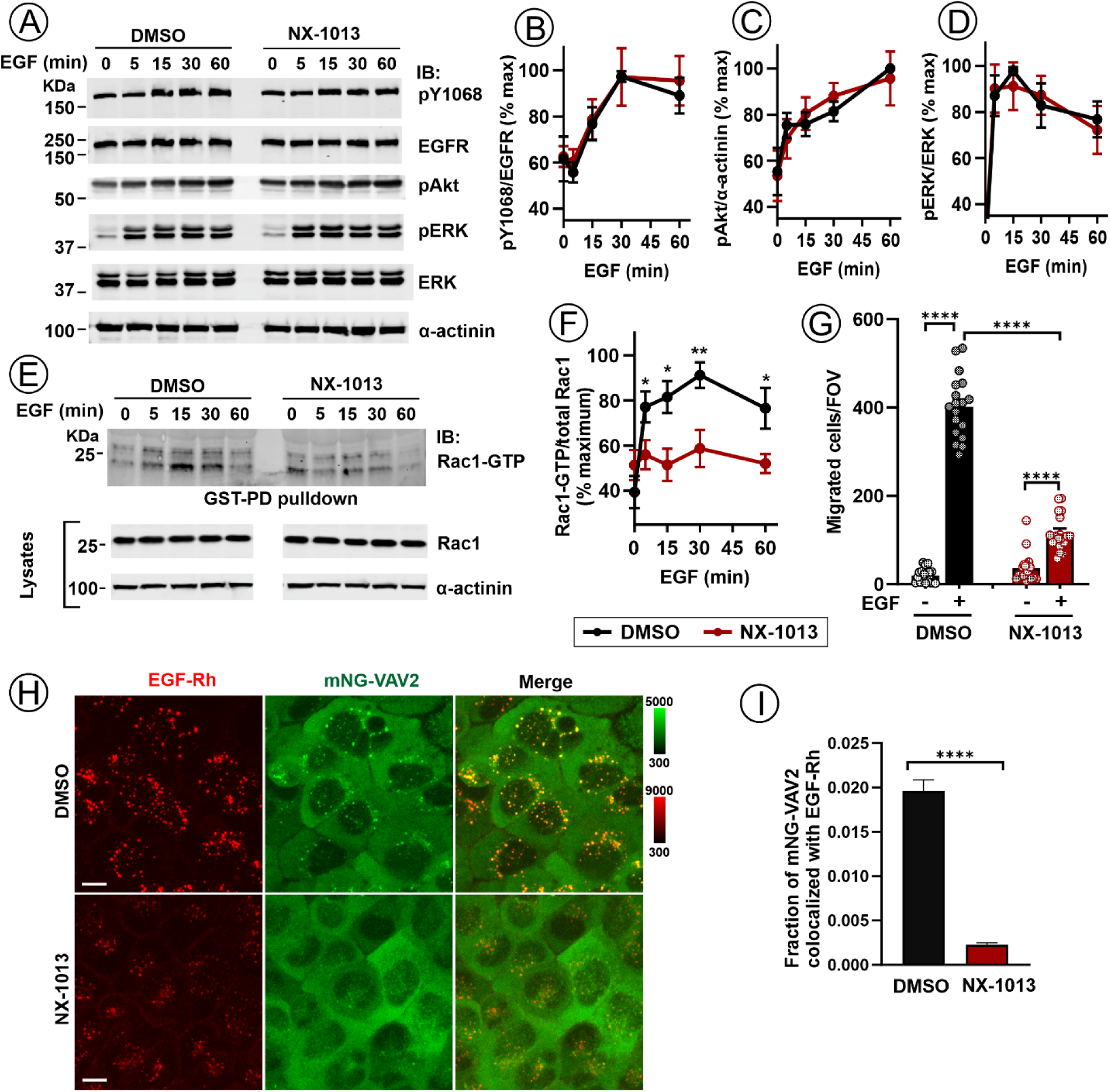
Effect of NX-1013 on EGFR signaling dynamics. (**A**) HSC3 cells were preincubated with DMSO or 100 nM NX-1013 for 1 hr and stimulated with 4 ng/ml EGF for indicated times at 37°C. Cell lysates were electrophoresed and probed by western blotting with antibodies to pY1068, EGFR, pERK1/2, ERK1/2, pAkt and α-actinin (loading control). Representative images are shown. (**B-D**) Quantification of phosphorylation of EGFR (pY1068) (***B***), Akt (***C***) and ERK1/2 (***D***) from western blots exemplified in ***A***. Graphs represent mean values (± SEM, n = 6 independent experiments) of the ratios pY1068/EGFR, pERK/ERK and pAkt/actinin immunoreactivities normalized to a maximal corresponding ratio in each individual time-course in DMSO-treated cells. **(E)** HSC3 cells were preincubated with DMSO or 100 nM NX-1013 for 1 hr and stimulated with 4 ng/ml EGF for indicated times at 37°C. Rac1-GTP was pulled down from cell lysates. Pulldowns and aliquots of lysates were electrophoresed and probed by western blotting using Rac1 and α-actinin (loading control) antibodies. Lysate images are from the same blot. Representative experiment is shown. **(F)** Quantification of the amount of Rac1-GTP from western blots exemplified in ***E***. Graph represents mean values of Rac1-GTP/total Rac1 ratio normalized to a maximum corresponding value in DMSO-treated cells within each individual time-course experiment (± SEM; n = 6 independent experiments). P values are calculated using unpaired two-tailed Student’s T-test comparing DMSO versus NX-1013 at the same time points. *p<0.05 (p = 0.049, 5 min; p = 0.016, 15 min; p = 0.040, 60 min); **p<0.01 (p = 0.0091). **(G)** Migration of HSC3 cells in Transwell chambers towards 4 ng/ml EGF gradient in the presence of DMSO or 100 nM NX-1013 was measured as described in ‘Methods’. Bars represent mean values (± SEM; n =3 independent experiments, 6 FOV each experiment) of numbers of cells migrated to the bottom surface of chamber. P values are calculated by one-way ANOVA. ****p<0.0001. **(H)** Localization of mNG-VAV2 in HSC3 cells was examined after stimulation of cells with 4 ng/ml EGF-Rh for 20 min by 3D live-cell imaging through 561 nm (EGF-Rh) and 515 nm (mNG-VAV2) channels. Maximum intensity projections of z-stacks are presented. Fluorescence intensity scales are identical for all images and shown on the right of images. Scale bars, 10 μm. **(I)** Quantifications of the fraction of mNG-VAV2 co-localized with EGF-Rh in images exemplified in (***H***). Bar graph represents mean values (± SEM; n=5 FOVs) of the fraction of mNG-VAV2 co-localized EGF-Rh. Unpaired two-tailed Student T-test between DMSO and NX-1013-treated cells was used to calculate p value. ****p<0.0001. Representative of 3 independent experiments.

In contrast with the signaling processes triggered from the plasma membrane, EGF-induced activation of Rac1, a GTPase involved in the regulation of cell motility, and EGFR-dependent cell migration are highly sensitive to the inhibition of CME and overexpression of CBLB mutants in HSC3 cells (26, 33). Therefore, we examined the dynamics of Rac1 activation in HSC3 cells stimulated with EGF in the presence of NX-1013. Measurements of Rac1 activity by pulldown of GTP-loaded Rac1 showed that NX-1013 significantly reduced this activity during the first hour of EGF (4 ng/ml) stimulation (Fig. 7E-F). We further tested whether NX-1013 affects HSC3 cell motility by performing Boyden chamber cell migration assays measuring cell migration in the presence of EGF concentration gradient. Figure 7G shows that NX-1013 dramatically decreased the number of migrating cells in the presence of EGF. To further examine the mechanisms underlying the effect of NX-1013 on Rac1 activation and cell migration, we examined the localization of VAV2, a major GDP/GTP exchange factor for Rac1 in HSC3 cells (33). To this end, gene-engineered HSC3 cells expressing endogenous VAV2 tagged with mNeonGreen fluorescent protein (mNG-VAV2) were stimulated with EGF-Rh. While mNG-VAV2 was highly co-localized with EGF-Rh in endosomes in control cells, mNG-VAV2 was virtually absent in endosomes of NX-1013-treated cells (Fig. 7H-I). CBL activity and tyrosine phosphorylation were demonstrated to be necessary for VAV2 and Rac1 activation in Abl-transformed fibroblasts (39). Therefore, it is possible that in addition to the reduced amount of EGFR in endosomes, NX-1013 affects the potential interaction of CBLs with VAV2, which may explain the dramatic reduction of VAV2 in endosomes containing active EGFR. Altogether, the data in Fig. 7E-I show that the endosomal localization of VAV2, Rac1 activation and EGF-dependent cell motility are highly sensitive to the activity of CBLs, and possibly to EGFR CME, which is consistent with the hypothesis that RTK-Rac1 signaling leading to cell migration is operational in endosomes (33, 40, 41).

To examine the long-term effects of NX-1013 on the EGFR activity in HSC3 cells under steady-state growth conditions, the cells were exposed to EGF for 6 and 24 hours. Measurements of total EGFR protein and active EGFR (pY1068 immunoreactivity) demonstrated that NX-1013 did not significantly affect the levels of total EGFR and active EGFR, although the amount of tyrosine phosphorylated EGFRs tended to be higher at the 6-hour time point in NX-1013-treated cells (p=0.055) (Figure 8A-D). As mentioned above, EGFR turnover and EGF-induced degradation of active and total EGFR are very slow in HSC3 cells due to the saturation of sorting pathways by the large amount of EGFR (26, 33). Therefore, as a proof of principle and to demonstrate that NX-1013 may influence long-term behavior of EGFR in cells with low levels of EGFR, similar experiments were performed in HeLa cells. In these cells, active and total EGFR were rapidly degraded in the presence of a sub-saturating EGF concentration (10 ng/ml) (Fig. 8E-H). NX-1013 substantially slowed down receptor turnover, which is especially evident for active EGFR (Fig. 8E-H). Strikingly, even in HeLa cells, non-ubiquitinated EGFR was degraded with a half-life time of ∼ 6 hrs in the presence of 10 ng/ml EGF (Figs. 8F and H). These kinetics are reminiscent of the slow degradation of a transmembrane cargo that is not ubiquitinated in experiments when the recycling is blocked (42). In the case of EGFR, it is possible that dimerization and oligomerization of EGFR minimize receptor recycling even in the absence of receptor ubiquitination, thus retaining a relatively large pool of EGFR in late endosomes which is eventually exposed to proteolytic enzymes. Such a mis-targeting was described for the traffic of the transferrin receptor bound to multivalent receptor-antibody complex (43).

**Figure 8.**
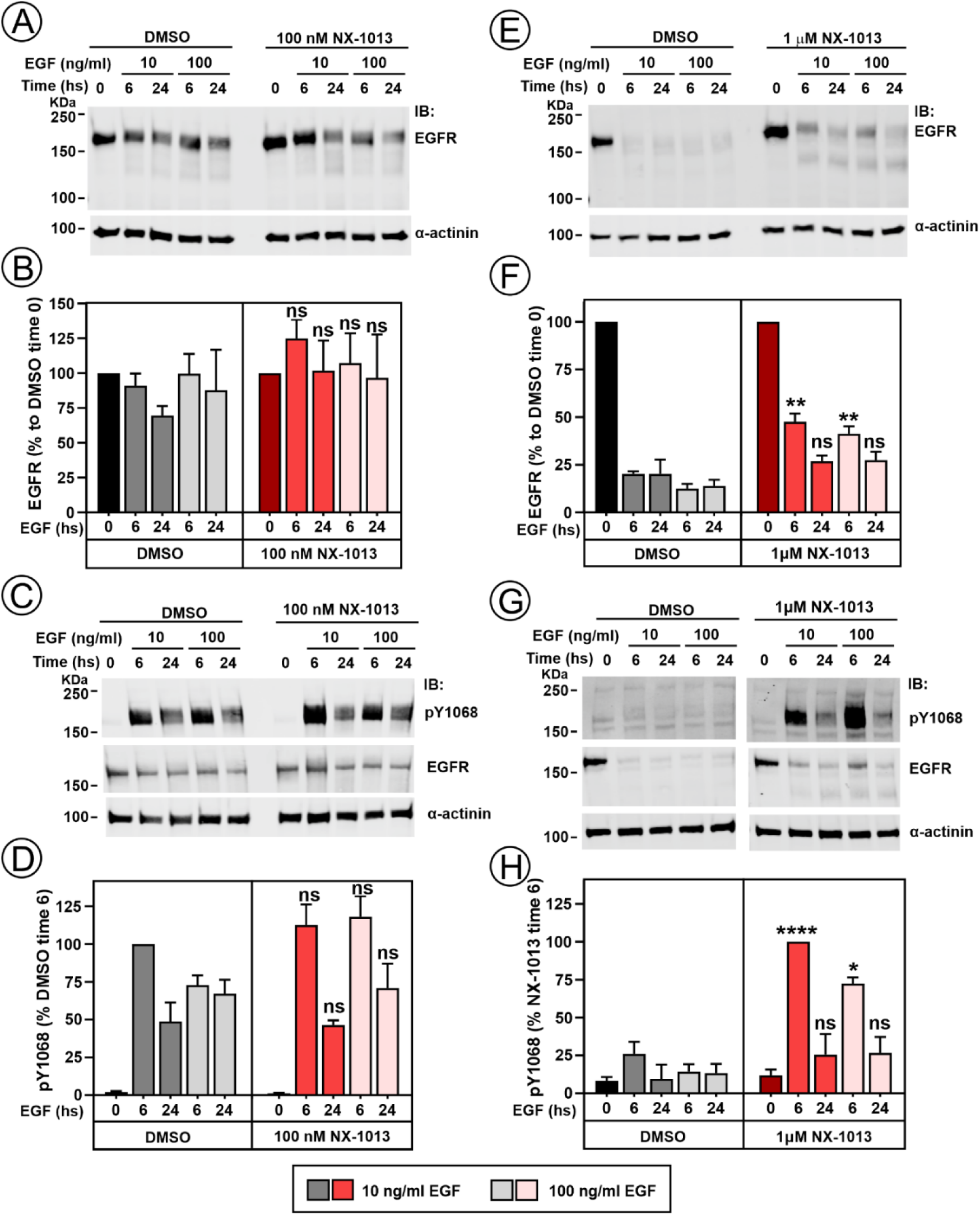
Long-term effect of NX-1013 on EGFR levels and activity. HSC3 (**A-D**) or HeLa cells (**E-H**) were preincubated with cycloheximide, and either DMSO or 100 nM NX-1013. The cells were then incubated with 10 or 100 ng/ml EGF in the same media for 6 hr or 24 hrs. Cells were lysed either in TGH lysis buffer in which orthovanadate and NEM were omitted (**A**, **B**, **E**, **F**) or in the complete TGH lysis buffer (**C**, **D**, **G**, **H**). Lysates were electrophoresed and probed by western blotting with antibodies to pY1068, EGFR, and α-actinin (loading control). Representative images are shown. Quantification of EGFR protein in **B** and **F** was performed by correcting the immunoreactivity of EGFR by the loading control and then normalizing the corrected values to the corrected value at time=0 in DMSO for each individual experiment. Quantification of EGFR phosphorylation (pY1068 signal) in **D** and **H** was performed by correcting the immunoreactivity of pY1068 by the loading control, and then normalizing the corrected values to the corrected value at time = 6 hs in DMSO in **D** and to the corrected value at time = 6 hrs in NX-1013 in **H** (because of the faint signal in DMSO-treated cells at time = 6 hrs) in each individual experiment. Bars represent mean values (± SEMs; n = 3 independent experiments). P values were calculated using unpaired two-tailed Student’s T-test comparing NX-1013 to DMSO for each time point. *p<0.05; **p<0.01; ****p<0.0001. See Table S2 for actual p values.

### Concluding remarks

Our study demonstrates that NX-1013 is highly effective in blocking E3 ligase activity of CBL proteins with a higher efficiency towards CBLB than CBL. In the EGFR system, inhibition of CBLB and CBL has a strong, although partial effect on the receptor endocytic trafficking. This illustrates a remarkable robustness of the endocytic machinery involved in ligand-induced down-regulation of EGFR with the considerable capacity to internalize and degrade non-ubiquitinated EGFR. Our characterization of the effects of NX-1013 on EGFR provides a paradigm for using NX-1013-like compounds for the quantitative analysis of the contribution of ubiquitination to the endocytic trafficking of other RTKs regulated by CBLs, such as c-MET, Axl, receptors for PDGF, VEGF and CSF, and others.

The robustness of EGFR endocytosis (existence of redundant endocytosis mechanisms) is one of the explanations for the lack of upregulating effects of NX-1013 on EGFR activity and signaling in EGFR-dependent carcinoma cells with high EGFR levels, such as HSC3 cells. Clinical trials with NX-1607 may provide important insights on whether inhibition of CBLs can elevate EGFR activity in cancers with low expression level of EGFR or in normal tissues. In addition to the robustness of EGFR endocytosis, an important consideration here is that concentrations of EGFR ligands in tumors, human plasma and interstitial fluids accessible to receptors in most tissues are extremely low, and therefore, the level of ligand-bound, active EGFR *in vivo* is correspondingly low [(44) and references therein]. Therefore, although NX-1013 caused a build-up of active EGFR in HeLa cells expressing low EGFR levels in the “proof-of-principle” experiments presented in Figure 8, EGF concentrations used in these experiments were significantly higher than the physiological concentrations of EGFR ligands (<1-2 ng/ml). The latter concentrations do not induce a significant long-term downregulation of EGFR in HeLa cells (34, 45), and therefore, a potential upregulating effect of the partial inhibition of EGFR endocytosis on its activity in cells with low/moderate levels of EGFR is expected to be minimal. Finally, given ubiquitous expression and relative abundance of CBL in most tissues, it is likely that a therapeutic dosage of NX-1607 series designed to inhibit CBLB would have only a moderate impact on CBL activity in non-hematopoietic cells. Therefore, while having upregulating effects on T and NK cell activities and their tumor infiltration (27), NX-1607 series compounds may only marginally affect EGFR ubiquitination, endocytosis and activity in most cell types, much like what we observed in HeLa cells when using 100 nM NX-1013 (Figs. 2E-F, S1 and 5). Thus, our studies support and highlight the promise of highly efficient small-molecule inhibitors of CBLB as effective cancer Immunotherapeutics.

## METHODS

### Chemistry

**–** The synthesis of NX-1013 is described in WO 2020/210508.

### Reagents

**-** Recombinant human EGF was from BD Biosciences (San Jose, CA). EGF-Rh was purchased from Invitrogen (Pittsburgh, PA). Actual concentration was determined in ^125^I-EGF competition experiments in comparison with unlabeled human EGF. Antibodies to EGFR (mRb #4267), pY1068 (Mo #2236), ERK1/2 (Mo #4696), pERK1/2 (Rb # 9101), pAkt (Mo #9272), α-actinin (Rb #6487), UbcH5c (Rb #4330) and CBL (Rb # 2179) were from Cell Signaling Technology (Danvers, MA). Mouse monoclonal antibodies to ubiquitin (P4D1, sc-8017), β-actin (sc-8432), CBLB (sc-8006) were from Santa Cruz Biotechnology (Dallas, TX). Antibodies for Rac1 (Mo, 610650) and EEA.1 (Mo, 610417) were from BD Transduction. EGFR monoclonal antibody 528 (Mab528) was from ATCC (Manassas, VA). The antibody to CHC (ab21679) and rabbit EEA1 (ab2900) were purchased from Abcam, to Grb2 (Rb PA5-27151) from Invitrogen and to LAMP1 (H4A3) from DSHB. The antibody to α-adaptin (AC1-M11, MA3-061) and Hoechst 33342 staining solution were from Thermofisher Scientific (Pittsburgh, PA). Other chemicals were from Fisher Scientific or Sigma if not indicated otherwise.

### Cell Culture

– Wild-type HSC3 cells and gene-edited HSC3/mNG-VAV2 cells generated previously (33) were maintained in DMEM supplemented with 5% fetal bovine serum (FBS). HeLa cells were maintained in DMEM supplemented with 10% FBS. HSC3 and HeLa cells were authenticated by STR profiling. All cells were mycoplasma-free. The cells were used for most experiments while growing in serum-containing media (not serum-starved) unless stated otherwise.

### RNAi interference

**–** We used CHC siRNA (duplex 2), μ2 subunit of AP-2 (duplex 4); [both - see Table S1 in(38)], Grb2 (36), CBL and CBLB (24), and UbcH5b/c (9). siRNA duplexes were resuspended in 1× siRNA universal buffer (Dharmacon, Inc., Lafayette, Colorado) to 20 μM prior to transfection. Non-targeting siRNA was from Qiagen. Cells grown in 35-mm dishes (60-80% confluency) were transfected with siRNA duplexes at a final concentration of 100 nM with DharmaFECT 1 reagent (Dharmacon) following manufacturer’s protocol. After 48 hrs, the cells were transfected with siRNAs again and re-plated to 12-well plates and used for experiments 2 days later.

### EGFR immunoprecipitation and western blotting

**-** The cells were stimulated with EGF at 37°C, washed with ice-cold Ca^2+^, Mg^2+^-free PBS and lysed in TGH buffer [1% Triton X-100, 10% (vol/vol) glycerol, 50 mM Hepes, 50 mM NaCl, 1 mM EDTA, 10 mM n-ethylmaleimide (NEM), 1 mM orthovanadate, and other phosphatase and protease inhibitors] as described in (26). Lysates were cleared by centrifugation and used for EGFR immunoprecipitation (1-1.5 mg protein) with the antibody 528 (10 μg/sample). The immunoprecipitates and aliquots of lysates (50 µg protein in HSC3 and 70 µg in HeLa) were resolved by SDS-PAGE (7.5%- or 10%-gels, respectively), followed by the transfer to the nitrocellulose membrane. Western blotting was performed by incubating with appropriate primary antibodies followed by secondary antibodies conjugated to far-red fluorescent dyes (IRDye-680 and -800) and detection using an Odyssey Li-COR system. Quantifications were performed using Li-COR software.

To examine long-term effects of NX-1013 on EGFR degradation, the cells were pre-incubated with 10 µM cycloheximide and NX-1013 (or DMSO) for 30 min in DMEM supplemented with 0.1% BSA. The cells were then incubated in DMEM containing 0.1% BSA with 0, 10 and 100 ng/ml EGF in the presence of 10 µM cycloheximide and NX-1031 (or DMSO) for 0, 6 and 24 hrs at 37°C. Cells were lysed either in complete TGH buffer or TGH in which NEM and orthovanadate were omitted. Pre-cleared lysates (50 µg protein in HSC3 and 70 µg in HeLa) were resolved by SDS-PAGE, transferred to the nitrocellulose membrane and probed by Western blotting as described above.

### Measurement of EGFR internalization rates using ^125^I-EGF

**-** Mouse receptor-grade EGF (Collaborative Research, Inc.) was iodinated as described (46). The cells were incubated with ^125^I-EGF (1-10 ng/mL) for 1-6 min at 37°C. The time–course of ^125^I-EGF internalization was used to calculate the specific internalization rate constant *k*_e_ as the linear regression coefficient of the dependence of the ratio of internalized/surface ^125^I-EGF on time as described (46). *k*_e_ values were also estimated using internalized/surface ratio values at a 6-min time point by dividing this ratio by 6, because of the amount of ligand at the cell surface and the linearity of the uptake was maintained during the first 6 min. The low concentration of ^125^I-EGF and short incubation times were used to avoid saturation of the CME pathway and minimize contribution of recycling, respectively. In another approach to examine EGFR internalization, cells were preincubated with ^125^I-EGF for 1 hr at 4°C to allow the equilibrium binding of ^125^I-EGF to cell-surface receptors in the absence of endocytosis and then chased at 37°C to allow endocytosis (Figure S3).

### Fluorescence microscopy and image analysis

– For live-cell imaging using spinning disk confocal system, the cells were grown on 35 mm Mat-Tek dishes, incubated with EGF-Rh at 37°C in phenol-free DMEM and immediately imaged at 37°C or room temperature (RT). For immunofluorescence labeling, the cells were grown on glass coverslips. After incubation with EGF-Rh, the cells were washed in PBS and fixed in freshly prepared 4% paraformaldehyde (PFA). Fixed cells were permeabilized with 0.1%Triton X-100 for 5 min, blocked in 3% BSA/PBS for 30 min at RT and incubated with Mab528, EEA1 or LAMP1 antibodies for 1 h at RT. Cells were then incubated with secondary antibodies for 1 h at RT. The nuclei were stained with Hoechst 33342 during incubation with secondary antibodies. Cells were mounted in ProLong Gold. To visualize cell-surface EGFR, the cells were fixed and stained with Mab528 without cell permeabilization.

Z-stack of 20-30 x-y confocal images were acquired using a Marianas spinning disk confocal imaging system based on Zeiss Axio Observer Z1 inverted fluorescence microscope system equipped with 63x Plan Apo PH NA1.4 objective, and controlled by SlideBook software (Intelligent Imaging Innovation, Denver, CO) as described (26). All image acquisition settings were identical for all experimental variants in each experiment.

To estimate the relative extent of colocalization of EGF-Rh or EGFR (Mab528) with EEA1 and LAMP1, 3D images were deconvolved using iterative constrain algorithm. A segment mask was then generated to select EEA1 or LAMP1 positive voxels detected through the 640 nm channel (EEA1 or LAMP1 masks) from background-subtracted images. Another segment mask was generated in the 561 nm or 488 nm channel to include total cellular EGF-Rh or EGFR fluorescence, respectively (“EGF” or “EGFR” mask, respectively). A “Colocalization” mask was then generated to select voxels overlapping in the EEA1 or LAMP1 masks with the EGF or EGFR mask. The sum fluorescence intensity through the 561 nm or 488 nm channel in the Colocalization mask was divided by the sum intensity of, correspondingly, the EGF or EGFR mask in each field of view (FOV) typically depicting 30-40 cells to calculate the fraction of EGF-Rh or EGFR colocalized with EEA1 or LAMP1 of the total cellular amount of EGF-Rh or EGFR. The intensity thresholds used to generate masks were identical for all variants in each experiment.

The estimate the fraction of cellular mNG-VAV2 (515 nm channel) in EGF-Rh-containing endosomes, a mask containing endosomal EGF-Rh (“EGF-Rh”; 561 nm channel) and a background mask were generated as described previously (47). The mean fluorescence intensity of mNG-VAV2 per voxel in the EGF-Rh mask was determined by subtracting the mean intensity of the background mask from that value in the EGF-Rh mask in each FOV. The sum of background-subtracted mNG-VAV2 fluorescence in the EGF-Rh mask was then calculated by multiplying the mean intensity by the voxel number of the EGF-Rh mask in each FOV and divided by the sum of total cellular mNG-VAV2 fluorescence per FOV to obtain the fraction of endosomal mNG-VAV2.

### Lattice LightSheet Microscopy

**–** Cells grown on 5-mm coverslips were placed in the imaging chamber of a Lattice LightSheet (LLS) V2 system (Intelligent Imaging Innovation, Inc., Denver, Colorado, USA). Excitation was achieved using 560 nm laser through an excitation objective (Special Optics 28.6x 0.7 NA 3.74-mm water immersion lens) and is detected via a Nikon CFI Apo LWD 25x 1.1 NA water immersion lens with a 2.5x tube lens using Hamamatsu Fusion BT sCMOS camera. Live cells were imaged in 3.0 ml of phenol-free DMEM at 37°C. 3D images (stacks of 101 images at 0.2 μm intervals) were acquired every 30 sec starting one minute after injection of EGF-Rh (10 ng/ml). Time-lapse images were deconvolved using the constrained iterative algorithm of SlideBook and presented as time-lapse sequences of maximum intensity projections of 3D images.

### Boyden chamber cell migration assays

**-** Boyden Chamber migration assays were carried out in sterile tissue culture plate inserts with polycarbonate membranes (VWR, Cat #10769-234). The membrane side facing the bottom of the 24-well plate was coated with fibronectin (50 µg/ml) before plating 10^5^ serum-starved cells in the top chamber of the inserts. The bottom chamber was filled with DMEM containing 0.1% BSA and 4 ng/ml EGF. Migration was allowed to proceed for 4 hrs at 37°C. Unattached cells from the top chamber were removed by suction with a cotton swab, and the cells remaining on the membrane and migrated to the bottom surface were fixed in 4%PFA for 15 min. Then, the inserts were washed twice in PBS, and cell nuclei were stained with Hoescht 33342 for 15 min. Finally, inserts were washed with excess water by immersion and allowed to air-dry in the dark prior imaging. Cells migrated to the bottom surface of the membrane were then counted using 10X objective from 8 random FOVs.

### Rac1-GTP GST-Pull Down Assay

- GST-fused Rac1·GTP binding fragment 70-117 aa of PAK1 (GST-PBD) in pGEX-2T was purchased from Addgene (#12217). Bacterially expressed GST-PBD was pulled down from precleared lysates using glutathione-Sepharose 4B beads (Cytiva, Sweden). The beads were washed in Ca^2+^, Mg^2+^-free PBS and immediately used in experiments. HSC3 cells were serum starved overnight and pre-incubated 1 hr with 100 nM NX-1013 or DMSO. Then, the cells were stimulated with 4 ng/ml EGF (in presence of NX-1013 or DMSO). The cells were washed with ice-cold DPBS and solubilized in 1% Triton X-100 lysis buffer (150 mM NaCl, 20 mM MgCl_2_, 50 mM Tris⋅HCl, pH 8, phosphatase and protease inhibitors). GST-PBD beads were incubated with 0.8-1.0 mg of precleared cell lysates for 1 h at 4°C. After extensive washes of beads with lysis buffer, proteins were eluted from beads in 2× sample buffer and heated for 5 min at 95°C. GST pulldowns and aliquots of cell lysates were resolved by electrophoresis, transferred to nitrocellulose membranes, and the membranes were probed by Western blotting as described above. Blots were imaged and analyzed using Odyssey Infrared Imaging system (LI-COR Biosciences).

### TR-FRET Probe Displacement Assay

– The binding affinity of NX-1013 to CBLB and CBL was assessed using TR-FRET probe displacement assays. The assays employed FL-436, a BODIPY-fluorescein-labeled analog of NX-1013, as the fluorescent probe. Assays were conducted in 384-well plates at room temperature in a final volume of 10 µl. Recombinant CBLB (0.125 nM) or CBL (0.5 nM) was pre-incubated for 1 hr with serial dilutions of NX-1013 (in 1% DMSO final concentration) in assay buffer containing 20 mM HEPES (pH 7.5), 150 mM NaCl, 0.01% Triton X-100, 0.01% BSA, and 0.5 mM TCEP. Following pre-incubation, the FL-436 probe was added at concentrations corresponding to its Kd values, along with streptavidin-terbium (Cisbio). Final concentrations were 200 nM FL-436 and 0.5 nM streptavidin-terbium for CBLB, and 425 nM FL-436 and 2 nM streptavidin-terbium for CBL. Plates were incubated for an additional hour before fluorescence measurements. TR-FRET signals were measured using an EnVision plate reader (PerkinElmer) with excitation at 340 nm and emission at 520 nm and 620 nm. The TR-FRET ratio (520/620 nm) was calculated, and background signal (in the absence of CBLB or CBL) was subtracted. Inhibitor activity was expressed as percent change in probe binding relative to DMSO controls. Dose-response curves were fitted using a standard Hill equation in GraphPad Prism (Y = B_min_ + ((B_max_ − B_min_) / (1+(IC_50_/X)^Hill))) to determine the IC_50_ values.

### Surface Plasmon Resonance (SPR) Binding Assay

**-** SPR experiments were performed using a Biacore S200 instrument (Cytiva Life Sciences). A Series S CM5 sensor chip was functionalized with NeutrAvidin (Invitrogen) via standard amine coupling. The sensor surface was activated with a 7 min injection of 195.5 mM EDC and 50 mM NHS, followed by immobilization of NeutrAvidin (125 μg/mL in 50 mM sodium acetate, pH 4.4) for 150 sec. Unreacted sites were quenched with 0.5 M ethanolamine (pH 8.0) for 3 min, resulting in a final NeutrAvidin density exceeding 10,000 resonance units (RU) per flow cell. All binding measurements were conducted at 25°C in assay buffer containing 20 mM HEPES (pH 7.5), 150 mM NaCl, 0.005% Tween-20, 1 mM TCEP, and 2% DMSO. Biotinylated CBLB and CBL proteins were diluted to 10–20 µg/ml in assay buffer and captured on the NeutrAvidin-coated surface to a final density of 2000–3000 RU. Flow cell 1 was used as a reference. To block residual biotin-binding sites, all flow cells were treated with two sequential 30 sec injections of 3 mM EZ-Link Amine-PEG2-Biotin (Thermo Scientific) at 30 µl/min. Test compounds were prepared in assay buffer and serially diluted in an 8-point, 3-fold dilution series. Binding was assessed using single-cycle kinetics with a 180 sec association phase at 50 µl/min, followed by a 7200 sec dissociation phase. Sensorgrams were corrected for DMSO content and double-referenced using blank injections and reference flow cell subtraction. Data were analyzed using Biacore Evaluation Software (Cytiva) and fit to a 1:1 binding model to derive kinetic parameters.

### Statistical data analysis

**-** Statistical analysis was performed using GraphPad Prism (La Jolla California USA). One-way ANOVA to control and unpaired T-test comparing each experimental group to control were normally used. Differences were considered significant if p-values are <0.05. P values and numbers of biological replicates are presented either in figure legends or in Tables S1-3 in Supplemental Materials (for Figures 2, 8, S1).

## Acknowledgements

We are thankful to Dr. Michael Lotze (University of Pittsburgh) for valuable discussion and advice, and to Brandon Bravo, Ketki Dhamnaskar and Stefan Gajewski (Nurix Therapeutics) for data collection or review. Medical illustration by Justin A Klein, CMI for Waterhouse Brands.

## Data sharing plan

All data generated and analyzed in this study are available from the corresponding author on reasonable request. This paper does not report original code. Any additional information required to reanalyze the data reported in this paper is available from the lead contact upon request.

## Supplemental Information

### Pinilla-Macua et al.

**Figure S1.**
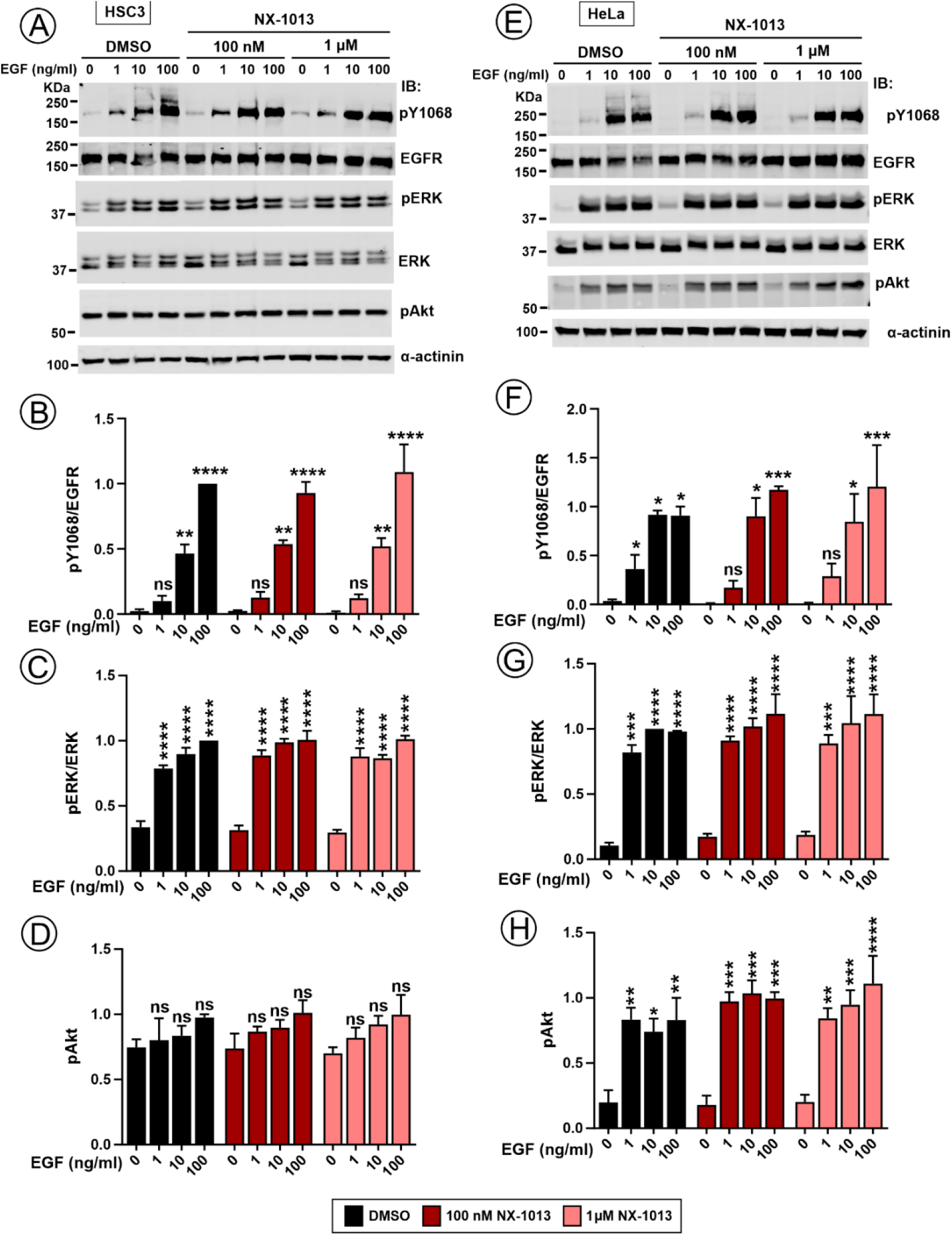
Effect of NX-1013 on EGFR phosphorylation and downstream signaling. **(A)** HSC3 cells were preincubated with DMSO, 100 nM or 1 μM NX-1013 for 1 hr and stimulated with 1-100 ng/ml EGF for 10 min at 37°C. Cell lysates were electrophoresed and probed by western blotting with antibodies to pY1068, EGFR, pERK1/2, ERK1/2, pAkt and α-actinin (loading control). Representative images are shown. (**B-D**) Quantification of phosphorylation of EGFR (pY1068) (***B***), ERK1/2 (***C***) and AKT (***D***) from western blots exemplified in ***A***. Bars represent mean values of phosphorylated proteins normalized to a corresponding maximum ratio in vehicle-treated cells in each individual experiment (± SEM; n = 3 independent experiments). P values for EGF-stimulated versus unstimulated cells were calculated using one-way ANOVA. **p<0.01; ****p<0.0001; ns, not significant (p>0.05). All differences for vehicle versus NX-1013 treated cells for same EGF-treatment variants were not significant (p>0.05). See Table S3 for actual p values. (**E**) HeLa cells were preincubated with DMSO, 100 nM or 1 μM NX-1013 for 1 hr and stimulated with 1-100 ng/ml EGF for 10 min at 37°C. Cell lysates were electrophoresed and probed by western blotting with antibodies to pY1068, EGFR, pERK1/2, ERK1/2, pAkt and α-actinin (loading control). Representative images are shown. (**F-H**) Quantification of phosphorylation of EGFR (pY1068) (***F***), ERK1/2 (***G***) and AKT (***H***) from western blots exemplified in ***E***. Bars represent mean values of protein phosphorylation proteins normalized to a corresponding maximum ratio in vehicle-treated cells in each individual experiment (± SEM; n = 3 independent experiments). P values for EGF-stimulated versus unstimulated cells were calculated using one-way ANOVA. *p<0.05; **p<0.01; ***p<0.001; ****p<0.0001; ns, not significant (p>0.05). All differences for vehicle versus NX-1013 treated cells for same EGF-treatment variants were not significant (p>0.05). See Table S3 for actual p values.

**Figure S2.**
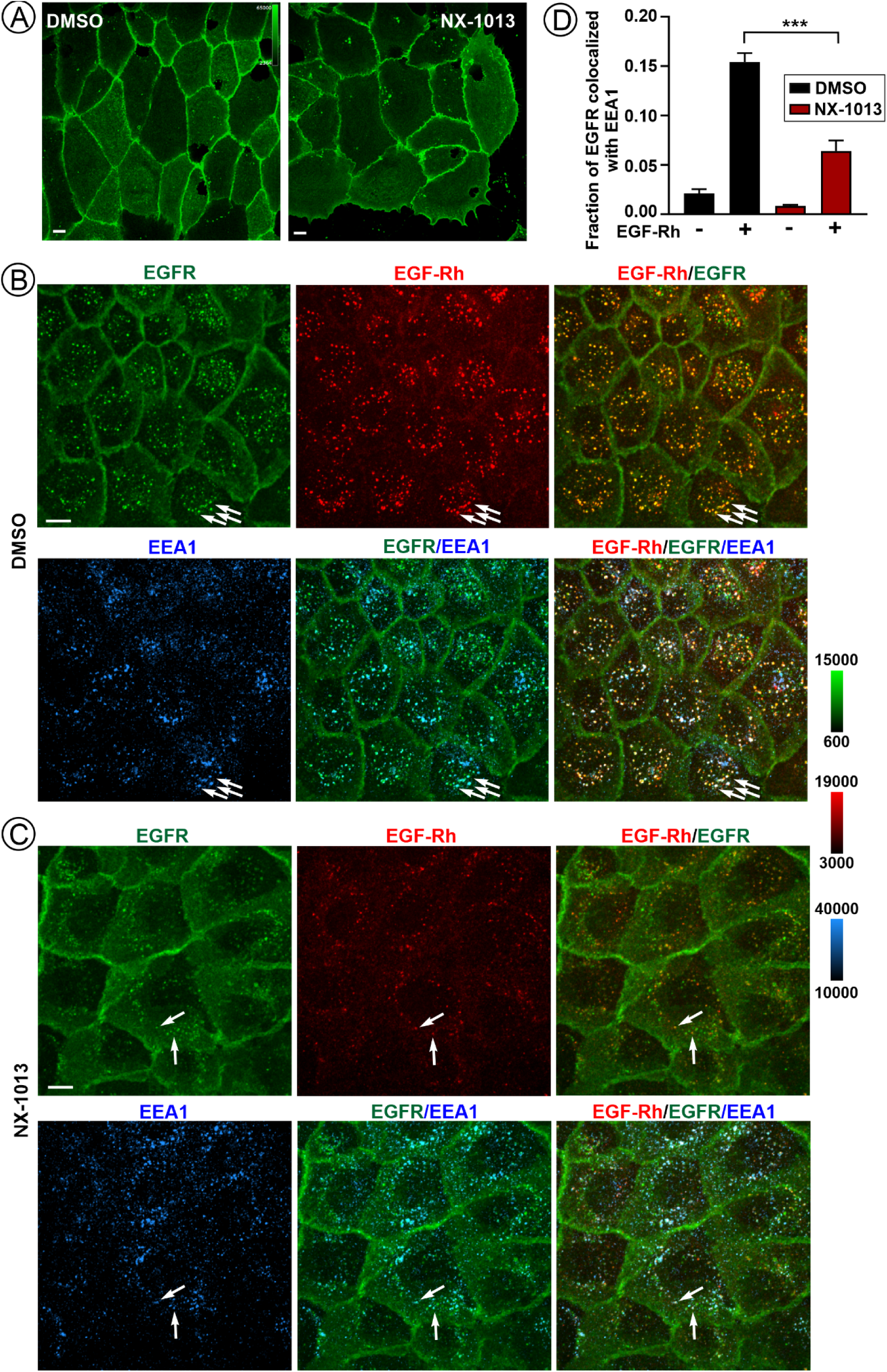
Effects of NX-1013 on EGFR expression, and internalization upon stimulation. Cells were pre-incubated with DMSO or 100 nM NX-1013 for 1 hr, fixed and labeled with EGFR Mab 528 without permeabilization **(A),** or incubated with 4 ng/ml EGF-Rh for 20 min at 37°C in the presence of DMSO **(B)** or NX-1013 **(C)**, fixed, permeabilized and immunolabeled with Mab528 and EEA1 antibodies. 3-D images were acquired through 640 nm (*blue*, EEA1), 561 nm (*red*, EGF-Rh) and 488 nm (*green*, EGFR) channels. Maximum intensity projections of z-stacks are presented. Fluorescence intensity scales are identical for all images (shown on the right). White arrows mark examples of EGFR puncta colocalized with EGF-Rh and EEA1. Scale bars, 10 µm. **(D)** Quantifications of the fraction of EGFR co-localized with EEA1 in images exemplified in ***B*** and ***C***. Bar graph represents mean values (± SEMs; n=5 FOVs) of the fraction of EGFR co-localized with EEA1. P values are calculated using unpaired two-tailed Student T-test comparing DMSO and NX-1013-treated cells. ***p=0.0002.

**Figure S3.**
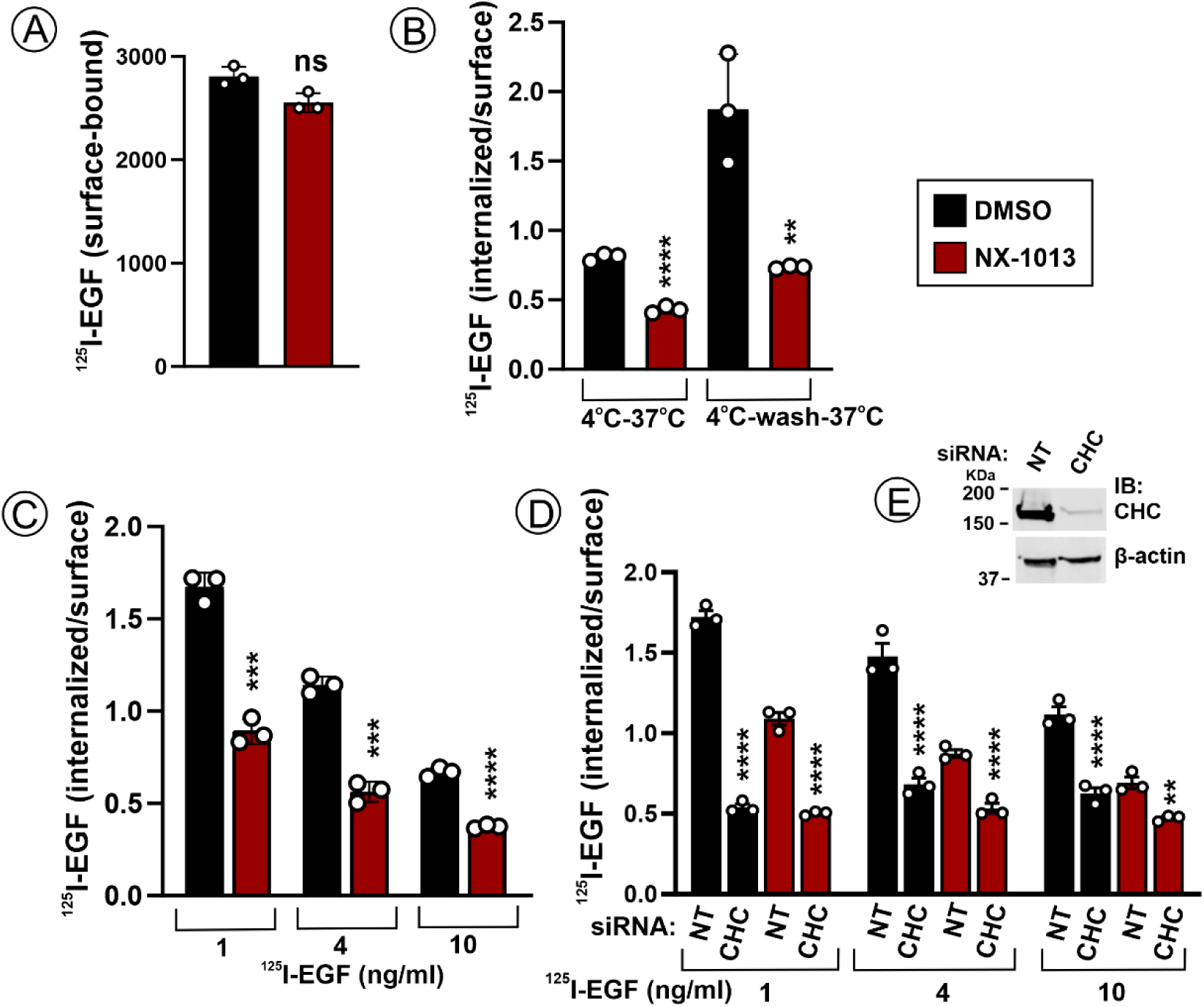
Effects of NX-1013 and CHC siRNA on internalization of 1-10 ng/ml ^125^I-EGF and after binding of ^125^I-EGF to cells at 4°C. **(A)** HSC3 cells were preincubated with DMSO or 100 nM NX-1013 for 1 hr, and then with ^125^I-EGF (1 ng/ml) for 45 min at 4°C. The amount of ^125^I-EGF associated with cells is shown in the bar graph. Mean values (± SEM; n=3) are shown. P values for NX-1013 versus DMSO were calculated using unpaired two-tailed Student’s T-test. ns, not significant. **(B)** HSC3 cells were preincubated with DMSO or 100 nm NX-1013 for 1 hr, and then with ^125^I-EGF (1 ng/ml) for 1 hour at 4°C, followed by the chase at 37°C for 6 min without cell wash or after washing off ^125^I-EGF. The ratio of internalized to surface ^125^I-EGF was measured as described in “Methods”. Bar graph represents mean values of internalized/surface ^125^I-EGF ratio (± SEM, n = 3). P values for NX-1013 versus DMSO were calculated as in ***A***. **p=0.0075; ****p<0.0001. **(C)** HSC3 cells were preincubated with DMSO or 100 nM NX-1013 for 1 hr, and then with ^125^I-EGF (1-10 ng/ml) for 6 min (1 and 4 ng/ml) or 5 min (10 ng/ml). The ratio of internalized to surface ^125^I-EGF was measured as described in “Methods”. Bar graph represents mean values of internalized/surface ^125^I-EGF ratio (± SEM, n = 3). P values for NX-1013 versus DMSO were calculated as in ***A***. ***p=0.00017 for 1 ng/ml; ***p=0.00014 for 4 ng/ml; ****p<0.0001. **(D)** HSC3 cells were transfected with non-targeting siRNA (NT) or siRNA targeting CHC. Four days after transfection, the cells were preincubated with DMSO or 100 nM NX-1013 for 1 hr and then incubated with ^125^I-EGF (1, 4 or 10 ng/ml) for 6 min. Bar graphs represent mean values of internalized/surface ^125^I-EGF ratio (± SEM, n = 3), P values for NX-1013 versus DMSO were calculated using one-way ANOVA. **p<0.01; **p<0.0027; ****p<0.0001. **(E)** Representative western blot to show the extent of CHC depletion in experiments presented in (***D***).

**Figure S4.**
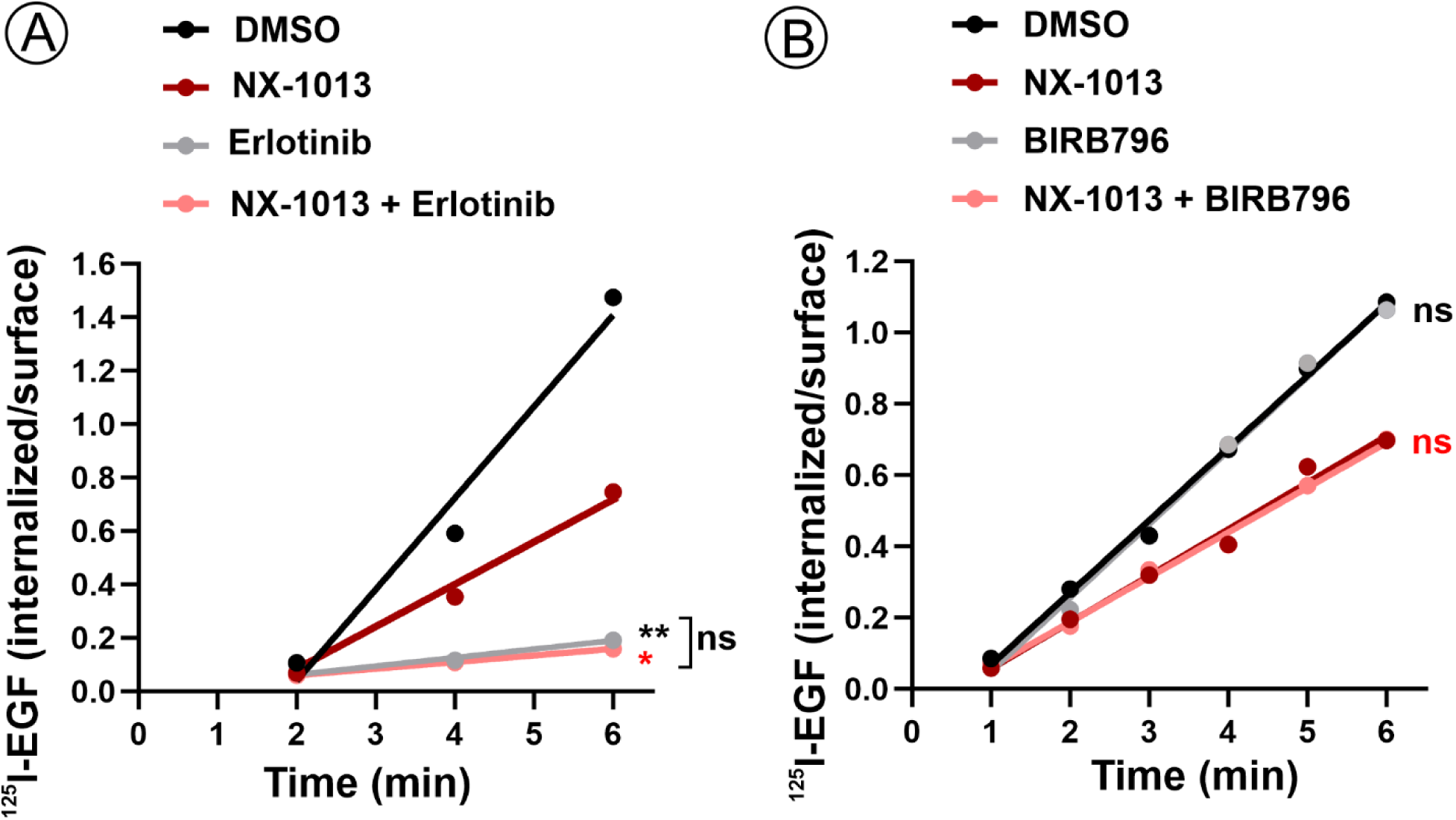
Effects of Erlotinib and BIRB796 on EGFR internalization. **(A)** HSC3 cells were preincubated with DMSO or 100 nM NX-1013 for 1 hr, and with or without Erlotinib (1 μM) and then with ^125^I-EGF (1 ng/ml) for 2-6 min. The ratio of internalized to surface ^125^I-EGF was measured as described in “Methods”. P values are calculated using one-way ANOVA. **p<0.0043 (p = 0.037 DMSO vs DMSO+Erlotinib; black asterisks); *p<0.05 (p = 0.049 NX-1013 vs Erlotinib + NX-1013; red asterisk). Difference between “Erlotinib” vs. “Erlotinib + NX-1013” is non-significant. **(B)** HSC3 cells were preincubated with DMSO or 100 nM BIRB796 for 3 hours, and with or without 100 nm NX-1013 for the last 1 hr of preincubation. The cells were then incubated with ^125^I-EGF (1 ng/ml) for 1-6 min. The ratio of internalized to surface ^125^I-EGF was measured as described in “Methods”. P values are calculated using one-way ANOVA. Differences between “DMSO” vs “BIRB796” or “NX-1013” vs “BIRB796 + NX-1013” are non- significant.

**Table S1:** Adjusted p values between each group and data sets in Figure 2.

| Data set | Multiple comparison test | Fig.2B | Fig.2C | Fig.2D | Fig.2F | Fig.2G |
| --- | --- | --- | --- | --- | --- | --- |
| DMSO | 0ng EGF vs.1 ng EGF | 0.0400 | 0.0053 | n.s. | n.s. | n.s. |
|  | 0ng EGF vs.10 ng EGF | 0.0042 | 0.0029 | 0.0057 | 0.0028 | 0.0254 |
|  | 0ng EGF vs.100 ng EGF | 0.0013 | 0.0489 | 0.0047 | 0.0068 | 0.0395 |
| NX-1013<br>100nM | 0ng EGF vs.1 ng EGF | n.s. | n.s. | n.s. | n.s. | n.s. |
|  | 0ng EGF vs.10 ng EGF | n.s. | 0.0323 | 0.0455 | 0.0144 | 0.0402 |
|  | 0ng EGF vs.100 ng EGF | n.s. | n.s. | 0.0172 | 0.0039 | 0.0011 |
| NX-1013 1μM | 0ng EGF vs.1 ng EGF | n.s. | n.s. | n.s. | n.s. | n.s. |
|  | 0ng EGF vs.10 ng EGF | n.s. | n.s. | n.s. | n.s. | n.s. |
|  | 0ng EGF vs.100 ng EGF | n.s. | n.s. | 0.0280 | n.s. | n.s. |
| 0ng EGF | DMSO vs. NX1013 100nM | n.s. | n.s. | n.s. | n.s. | n.s. |
| | DMSO vs. NX1013 1μM | 0.0320 | n.s. | n.s. | n.s. | n.s. |
| 1ng EGF | DMSO vs. NX1013 100nM | 0.0073 | 0.0237 | n.s. | n.s. | n.s. |
| | DMSO vs. NX1013 1μM | 0.0027 | 0.0100 | n.s. | n.s. | n.s. |
| 10ng EGF | DMSO vs. NX1013 100nM | <0.0001 | n.s. | n.s. | 0.0028 | n.s. |
| | DMSO vs. NX1013 1μM | <0.0001 | n.s. | n.s. | 0.0011 | n.s. |
| 100ng EGF | DMSO vs. NX1013 100nM | 0.0019 | n.s. | n.s. | 0.0072 | n.s. |
| | DMSO vs. NX1013 1μM | 0.0018 | n.s. | n.s. | 0.0026 | n.s. |

**Table S2:** Adjusted p values between each group and data sets in Figure 8.

| Multiple comparison test | Data set | Fig.8B | Fig.8D | Fig.8F | Fig.8H |
| --- | --- | --- | --- | --- | --- |
| DMSO vs.NRX-1013 | 0h EGF | n.s. | n.s. | n.s. | n.s. |
|  | 6h EGF 10 ng/ml | n.s. | n.s. | 0.0040 | <0.0001 |
|  | 24h EGF 10 ng/ml | n.s. | n.s. | n.s. | n.s. |
|  | 6h EGF 100 ng/ml | n.s. | n.s. | 0.0033 | 0.0017 |
|  | 24h EGF 100 ng/ml | n.s. | n.s. | n.s. | n.s. |

**Table S3:** Adjusted p values between each group and data sets in Figure S1.

| Data set | Multiple comparison test | Fig.S1 B | Fig. S1C | Fig. S1D | Fig. S1F | Fig. S1G | Fig. S1H |
| --- | --- | --- | --- | --- | --- | --- | --- |
| DMSO | 0ng EGF vs.1 ng EGF | n.s. | <0.0001 | n.s. | n.s. | 0.0001 | 0.0040 |
|  | 0ng EGF vs.10 ng EGF | 0.0127 | <0.0001 | n.s. | 0.0147 | <0.0001 | 0.0176 |
|  | 0ng EGF vs.100 ng EGF | <0.0001 | <0.0001 | n.s. | 0.0159 | <0.0001 | 0.0042 |
| NX-1013 100nM | 0ng EGF vs.1 ng EGF | n.s. | <0.0001 | n.s. | n.s. | <0.0001 | 0.0003 |
|  | 0ng EGF vs.10 ng EGF | 0.0025 | <0.0001 | n.s. | 0.0324 | <0.0001 | 0.0001 |
|  | 0ng EGF vs.100 ng EGF | <0.0001 | <0.0001 | n.s. | 0.0010 | <0.0001 | 0.0002 |
| NX-1013 1µM | 0ng EGF vs.1 ng EGF | n.s. | <0.0001 | n.s. | n.s. | <0.0001 | 0.0035 |
|  | 0ng EGF vs.10 ng EGF | 0.0037 | <0.0001 | n.s. | 0.0226 | <0.0001 | 0.0007 |
|  | 0ng EGF vs.100 ng EGF | <0.0001 | <0.0001 | n.s. | 0.0007 | <0.0001 | <0.0001 |
| 0ng EGF | DMSO vs. NX1013 100nM | n.s. | n.s. | n.s. | n.s. | n.s. | n.s. |
|  | DMSO vs. NX1013 1µM | n.s. | n.s. | n.s. | n.s. | n.s. | n.s. |
| 1ng EGF | DMSO vs. NX1013 100nM | n.s. | n.s. | n.s. | n.s. | n.s. | n.s. |
|  | DMSO vs. NX1013 1µM | n.s. | n.s. | n.s. | n.s. | n.s. | n.s. |
| 10ng EGF | DMSO vs. NX1013 100nM | n.s. | n.s. | n.s. | n.s. | n.s. | n.s. |
|  | DMSO vs. NX1013 1µM | n.s. | n.s. | n.s. | n.s. | n.s. | n.s. |
| 100ng EGF | DMSO vs. NX1013 100nM | n.s. | n.s. | n.s. | n.s. | n.s. | n.s. |
|  | DMSO vs. NX1013 1µM | n.s. | n.s. | n.s. | n.s. | n.s. | n.s. |

## LEGENDS TO VIDEOS

### VideoS1. Time-lapse imaging of EGF-Rh endocytosis in vehicle-treated HSC3 cells

Cells were pre-incubated with DMSO for 1 hour and then 3D images were acquired every 30 seconds starting 1 min after injection of EGF-Rh (10 ng/ml) for 30 min at 37°C using the LLS system. Maximum intensity projections of z-stack of x-y images are shown. Scale bar, 10 μm.

### VideoS2. Time-lapse imaging of EGF-Rh endocytosis in HSC3 cells treated with NX-1013

Cells were pre-incubated with 100 nM NX-1013 and then 3D images were acquired every 30 seconds starting 1 min after injection of EGF-Rh (10 ng/ml) in the presence of NX-1013 for 30 min at 37°C using the LLS system. Maximum intensity projections of z-stack of x-y images are shown. Scale bar, 10 μm.

## REFERENCES

1. M. Sibilia et al., The epidermal growth factor receptor: from development to tumorigenesis. Differentiation 75, 770–787 (2007).

2. S. Pastore, F. Mascia, V. Mariani, G. Girolomoni, The epidermal growth factor receptor system in skin repair and inflammation. J Invest Dermatol 128, 1365–1374 (2008).

3. M. L. Uribe, I. Marrocco, Y. Yarden, EGFR in Cancer: Signaling Mechanisms, Drugs, and Acquired Resistance. Cancers (Basel) 13 (2021).

4. M. A. Lemmon, J. Schlessinger, Cell signaling by receptor tyrosine kinases. Cell 141, 1117–1134 (2010).

5. A. Sorkin, L. K. Goh, Endocytosis and intracellular traficking of ErbBs. Exp Cell Res 315, 683–696 (2009).

6. S. Sigismund, L. Lanzetti, G. Scita, P. P. Di Fiore, Endocytosis in the context-dependent regulation of individual and collective cell properties. Nat Rev Mol Cell Biol 10.1038/s41580-021-00375-5 (2021).

7. M. von Zastrow, A. Sorkin, Mechanisms for Regulating and Organizing Receptor Signaling by Endocytosis. Annu Rev Biochem 10.1146/annurev-biochem-081820-092427 (2021).

8. L. K. Goh, A. Sorkin, Endocytosis of receptor tyrosine kinases. Cold Spring Harb Perspect Biol 5, a017459 (2013).

9. A. Fortian et al., Endocytosis of Ubiquitylation-Deficient EGFR Mutants via Clathrin-Coated Pits is Mediated by Ubiquitylation. Traffic 16, 1137–1154 (2015).

10. L. K. Goh, F. Huang, W. Kim, S. Gygi, A. Sorkin, Multiple mechanisms collectively regulate clathrin-mediated endocytosis of the epidermal growth factor receptor. J Cell Biol 189, 871–883 (2010).

11. S. Sigismund et al., Threshold-controlled ubiquitination of the EGFR directs receptor fate. The EMBO journal 32, 2140–2157 (2013).

12. R. Pascolutti et al., Molecularly Distinct Clathrin-Coated Pits Diferentially Impact EGFR Fate and Signaling. Cell Rep 27, 3049–3061.e3046 (2019).

13. G. Ahmad et al., Cbl-family ubiquitin ligases and their recruitment of CIN85 are largely dispensable for epidermal growth factor receptor endocytosis. The international journal o biochemistry & cell biology 57, 123–134 (2014).

14. F. Huang et al., Lysine 63-linked polyubiquitination is required for EGF receptor degradation. Proc Natl Acad Sci U S A 110, 15722–15727 (2013).

15. G. Levkowitz et al., Ubiquitin ligase activity and tyrosine phosphorylation underlie suppression of growth factor signaling by c-Cbl/Sli-1 [In Process Citation]. Mol Cell 4, 1029–1040 (1999).

16. C. H. Shen et al., ZNRF1 Mediates Epidermal Growth Factor Receptor Ubiquitination to Control Receptor Lysosomal Traficking and Degradation. Front Cell Dev Biol 9, 642625 (2021).

17. M. H. Schmidt, I. Dikic, The Cbl interactome and its functions. Nat Rev Mol Cell Biol 6, 907–918 (2005).

18. N. Rao, I. Dodge, H. Band, The Cbl family of ubiquitin ligases: critical negative regulators of tyrosine kinase signaling in the immune system. J Leukoc Biol 71, 753–763 (2002).

19. E. K. Grifiths et al., Cbl-3-Deficient Mice Exhibit Normal Epithelial Development. Molecular and Cellular Biology 23, 7708–7718 (2003).

20. P. Peschard, N. Ishiyama, T. Lin, S. Lipkowitz, M. Park, A conserved DpYR motif in the juxtamembrane domain of the Met receptor family forms an atypical c-Cbl/Cbl-b tyrosine kinase binding domain binding site required for suppression of oncogenic activation. J Biol Chem 279, 29565–29571 (2004).

21. G. Swaminathan, A. Y. Tsygankov, The Cbl family proteins: ring leaders in regulation of cell signaling. J Cell Physiol 209, 21–43 (2006).

22. L. Duan et al., Cbl-mediated ubiquitinylation is required for lysosomal sorting of epidermal growth factor receptor but is dispensable for endocytosis. J Biol Chem 278, 28950–28960 (2003).

23. B. L. M. Crotchett, B. P. Ceresa, Knockout of c-Cbl slows EGFR endocytic traficking and enhances EGFR signaling despite incompletely blocking receptor ubiquitylation. Pharmacol Res Perspect 9, e00756 (2021).

24. F. Huang, D. Kirkpatrick, X. Jiang, S. Gygi, A. Sorkin, Diferential regulation of EGF receptor internalization and degradation by multiubiquitination within the kinase domain. Mol Cell 21, 737–748 (2006).

25. S. Pennock, Z. Wang, A tale of two Cbls: interplay of c-Cbl and Cbl-b in epidermal growth factor receptor downregulation. Mol Cell Biol 28, 3020–3037 (2008).

26. I. Pinilla-Macua, A. Sorkin, Cbl and Cbl-b independently regulate EGFR through distinct receptor interaction modes. Mol Biol Cell 34, ar134 (2023).

27. S. Gajewski et al., Discovery and characterization of small molecule inhibitors of CBL-B that act as intramolecular glue to enhance T-cell anti-tumor activity. bioRxiv 10.64898/2026.01.22.701125, 2026.2001.2022.701125 (2026).

28. M. Galliotta, Gosling, J., Tenn-McClellan, A., Ranucci, S., Gomez Romo, J., Cohen, F., Hansen, G., Sands, A., Guiducci, C., Rountree, R., 824 NX-1607, a small molecule inhibitor of the CBL-B E3 ubiquitin ligase, promotes T and NK cell activation and enhances NK-mediated ADCC in a mouse lymphoma tumor model. Journal or ImmunoTherapy o Cancer 10 (2022).

29. G. P. Collins et al., A First-in-Human Phase 1 Trial of NX-1607, a First-in-Class Oral CBL-B Inhibitor, in Patients with Advanced Malignancies Including DLBCL. Blood 142, 3093–3093 (2023).

30. Y. Kobashigawa et al., Autoinhibition and phosphorylation-induced activation mechanisms of human cancer and autoimmune disease-related E3 protein Cbl-b. Proc Natl Acad Sci U S A 108, 20579–20584 (2011).

31. H. Dou et al., Structural basis for autoinhibition and phosphorylation-dependent activation of c-Cbl. Nat Struct Mol Biol 19, 184–192 (2012).

32. Y. Ohnishi, O. Lieger, M. Attygalla, T. Iizuka, K. Kakudo, Efects of epidermal growth factor on the invasion activity of the oral cancer cell lines HSC3 and SAS. Oral Oncol 44, 1155–1159 (2008).

33. I. Pinilla-Macua, S. Surve, A. Sorkin, Cell migration signaling through the EGFR-VAV2-Rac1 pathway is sustained in endosomes. J Cell Sci 138 (2025).

34. M. Perez Verdaguer et al., Mechanism of p38 MAPK-induced EGFR endocytosis and its crosstalk with ligand-induced pathways. J Cell Biol 220 (2021).

35. L. K. Opresko et al., Endocytosis and lysosomal targeting of epidermal growth factor receptors are mediated by distinct sequences independent of the tyrosine kinase domain. J. Biol. Chem. 270, 4325–4333 (1995).

36. X. Jiang, F. Huang, A. Marusyk, A. Sorkin, Grb2 Regulates Internalization of EGF Receptors through Clathrin-coated Pits. Mol Biol Cell 14, 858–870 (2003).

37. K. Umebayashi, H. Stenmark, T. Yoshimori, Ubc4/5 and c-Cbl Continue to Ubiquitinate EGF Receptor after Internalization to Facilitate Polyubiquitination and Degradation. Mol Biol Cell 19, 3454–3462 (2008).

38. F. Huang, A. Khvorova, W. Marshall, A. Sorkin, Analysis of clathrin-mediated endocytosis of epidermal growth factor receptor by RNA interference. J Biol Chem 279, 16657–16661 (2004).

39. A. M. Teckchandani, E. A. Feshchenko, A. Y. Tsygankov, c-Cbl facilitates fibronectin matrix production by v-Abl-transformed NIH3T3 cells via activation of small GTPases. Oncogene 20, 1739–1755 (2001).

40. A. Palamidessi et al., Endocytic traficking of Rac is required for the spatial restriction of signaling in cell migration. Cell 134, 135–147 (2008).

41. L. Menard, P. J. Parker, S. Kermorgant, Receptor tyrosine kinase c-Met controls the cytoskeleton from diferent endosomes via diferent pathways. Nature communications 5, 3907 (2014).

42. A. J. Evans, J. L. Daly, A. N. K. Anuar, B. Simonetti, P. J. Cullen, Acute inactivation of retromer and ESCPE-1 leads to time-resolved defects in endosomal cargo sorting. J Cell Sci 133 (2020).

43. C. R. Hopkins, I. S. Trowbridge, Internalization and processing of transferrin and the transferrin receptor in human carcinoma A431 cells. J Cell Biol 97, 508–521 (1983).

44. I. Pinilla-Macua, A. Grassart, U. Duvvuri, S. C. Watkins, A. Sorkin, EGF receptor signaling, phosphorylation, ubiquitylation and endocytosis in tumors in vivo. Eli e 6 (2017).

45. S. Sigismund et al., Clathrin-mediated internalization is essential for sustained EGFR signaling but dispensable for degradation. Dev Cell 15, 209–219 (2008).

46. A. Sorkin, J. E. Duex, Quantitative analysis of endocytosis and turnover of epidermal growth factor (EGF) and EGF receptor. Curr Protoc Cell Biol **Chapter** 15, Unit 15 14 (2010).

47. S. V. Surve et al., Localization dynamics of endogenous fluorescently labeled RAF1 in EGF-stimulated cells. Mol Biol Cell 30, 506–523 (2019).

